# Brain-wide mapping of proglucagon expression in mice identifies fasting-responsive GLP-1 neurons in the posterior hypothalamic nucleus

**DOI:** 10.64898/2026.08.10.743428

**Authors:** Gábor Wittmann, Andrea Kádár, Petra Mohácsik, Morten Grønbech Rasch, Yvette Ruska, István Várkonyi, Beáta Dorogházi, András Horváth, Zsolt Liposits, Balázs Gereben, Csaba Fekete

## Abstract

**Objective:** Glucagon-like peptide-1 (GLP-1), a peptide neurotransmitter in the brain, is synthesized from proglucagon, encoded by the glucagon gene (*Gcg*). Besides medullary GLP-1 neurons, *Gcg*-expressing neuron populations were identified in the olfactory bulb and basolateral amygdala. However, several lines of evidence suggest that additional *Gcg* neuron populations might exist.

**Methods:** We conducted a brain-wide mapping of *Gcg*-expressing cells by fluorescent *in situ* hybridization in C57BL/6J and FVB/Ant mice. Proglucagon and GLP-1 expression were studied with immunofluorescence. We characterized a *Gcg*-Cre;tdTomato mouse line and studied the expression of proglucagon-processing enzymes in *Gcg*-expressing neuron populations. We used adeno-associated virus-mediated tracing in *Gcg*-Cre mice to map the projections of hypothalamic *Gcg* neurons.

**Results:** *Gcg*-expressing neuron populations were identified in the olfactory bulb, claustrum, piriform cortex, basolateral amygdala, posterior hippocampus, posterior hypothalamic nucleus (PH), periaqueductal gray/dorsal raphe, and dorsal nucleus of the lateral lemniscus. These neurons express lower *Gcg* mRNA levels than medullary GLP-1 neurons. Proglucagon and GLP-1-immunoreactivity (C-terminus) were detected in almost all *Gcg*-expressing neuron populations, along with the mRNAs for prohormone convertases 1/3 and 2, enzymes generating GLP-1 or glucagon, respectively. Fasting markedly increased *Gcg* mRNA, proglucagon and GLP-1 synthesis in the PH. PH *Gcg* neurons project densely to the ventral and intermediate lateral septum, preoptic region, ventrolateral preoptic nucleus, lateral hypothalamus and zona incerta, establishing close contacts with both GLP-1 receptor-positive and -negative neurons.

**Conclusions:** Proglucagon is expressed in 9 distinct neuron populations. Feeding status regulates GLP-1 synthesis in PH neurons that likely control feeding- or energy balance-related functions.

## 1. Introduction

The glucagon-like peptide 1 (GLP-1) /GLP-1 receptor (GLP-1R) system of the brain has been the subject of intense research that has gained even more momentum since GLP-1 analogues have become widely used to treat type 2 diabetes and obesity. GLP-1 is generated by enzymatic processing from proglucagon (PG), encoded by the glucagon gene (*Gcg*) [1]. *Gcg* mRNA-expressing neuron populations have been identified in only three regions of the adult mammalian brain. The cells commonly referred to as GLP-1 neurons are located in the medulla, in the nucleus of the solitary tract (NTS) and the intermediate reticular nucleus (IRN) [2–8]. These cells were discovered soon after the first reports on glucagon-immunoreactive materials in the brain [9–12], initially by immunohistochemistry [2; 3], and subsequently by *in situ* hybridization (ISH) [4]. These neurons regulate satiety, feeding behavior and autonomic functions *via* widespread projections through the brain and spinal cord [13–15]. Another *Gcg*-expressing neuron population was discovered later by ISH in the glomerular layer of the rat olfactory bulb (OB) [8]. Recently, *Gcg* mRNA was also detected in the granule cell layer of the rat OB by RNAscope ISH [16], corroborating earlier observations of *Gcg* promoter-driven transgene expression in mice [17; 18]. Recent studies suggest a role for these cells in pancreatic insulin release [19; 20]. Finally, Zheng and colleagues [16] identified *Gcg*-expressing neurons in the basolateral amygdala (BLA). They confirmed active transcription of the *Gcg* locus in the BLA of *Gcg*-Cre rats by viral transduction, and pinpointed *Gcg* mRNA-expressing cells in the mouse BLA in the Allen Brain Atlas [21].

Several studies raised the possibility, however, that *Gcg* may be expressed by additional neuron populations, although at low levels, and possibly in a physiological state-dependent manner. *Gcg* mRNA was identified in the rat hypothalamus by Northern blot, although at 100-fold lower levels than in the brainstem [22]. *Gcg* mRNA was also detected by RT-PCR in the cortex, hypothalamus, cerebellum and hippocampus of adult mice and/or rats [23; 24]. Neurons immunoreactive for the PG-derived peptide, glicentin, were observed in the mediobasal hypothalamus of fasted, but not fed rats [25]. Transgenic mouse studies also mention scattered neurons expressing *Gcg* promoter-driven transgenes in various parts of the central nervous system [6; 16; 26; 27]. Although ectopic transgene expression may occur in these mice generated with early transgenic [6] or bacterial artificial chromosome (BAC) technology [27–29], we recently confirmed the presence of *Gcg* mRNA-expressing neurons in the mouse spinal cord [30], matching the distribution of a *Gcg* promoter driven transgene [31]. In addition, a large number of neurons expressing the Cre-dependent reporter were observed in the piriform cortex of *Gcg*-Cre/tdTomato rats generated by knock-in technology [16].

To search for additional *Gcg*-expressing neuron populations, we first conducted a brain-wide fluorescent ISH (FISH) study in fed and fasted mice. Having identified such populations, we used immunofluorescence (IF) to examine whether these cells synthesize PG and GLP-1. To facilitate the investigation of the newly recognized *Gcg* neuron groups, we thoroughly characterized a *Gcg*-Cre mouse line, and studied the expression of PG-processing enzymes in *Gcg*-expressing neuron populations. To corroborate that fasting-responsive hypothalamic *Gcg* neurons signal via GLP-1, we mapped their axonal projections by viral tract tracing and examined GLP-1R expression in their downstream neurons.

## 2. Results

### 2.1 Brain-wide distribution of Gcg-expressing cells in ad libitum fed mice

To detect *Gcg* mRNA, we performed FISH on serial coronal sections (∼200 µm apart) through the mouse brain, from the OB to the lower medulla. To identify potential sex- or strain-related differences, we examined male C57BL/6J (n=3; 10, 15 and 20 weeks /wks/ old), female C57BL/6J (n=3; 10-11 wks), and male FVB/Ant mice (n=3, 16 wks). *Gcg* expression pattern was very similar among all mice examined. We identified 9 *Gcg*-expressing neuron populations that were present in all mice, and one minor cell group, which was observed only in male C57BL/6J mice. *Gcg* expression was uniquely high in NTS/IRN neurons. All other populations expressed *Gcg* mRNA at much lower levels. The FISH signal completely filled the cytoplasm in NTS/IRN neurons, while in the other populations it generally appeared as distinct dots, merging only in occasional cells (**Figs 1-2**). To estimate the relative cell numbers of *Gcg*-expressing neuron populations, we counted *Gcg*-positive cells in male C57BL/6J mice (**Fig 2F**; n≥3 per region). We generally considered a cell *Gcg*-positive if it contained at least 3 hybridization dots (see section *4.13*).

**Figure 1.**
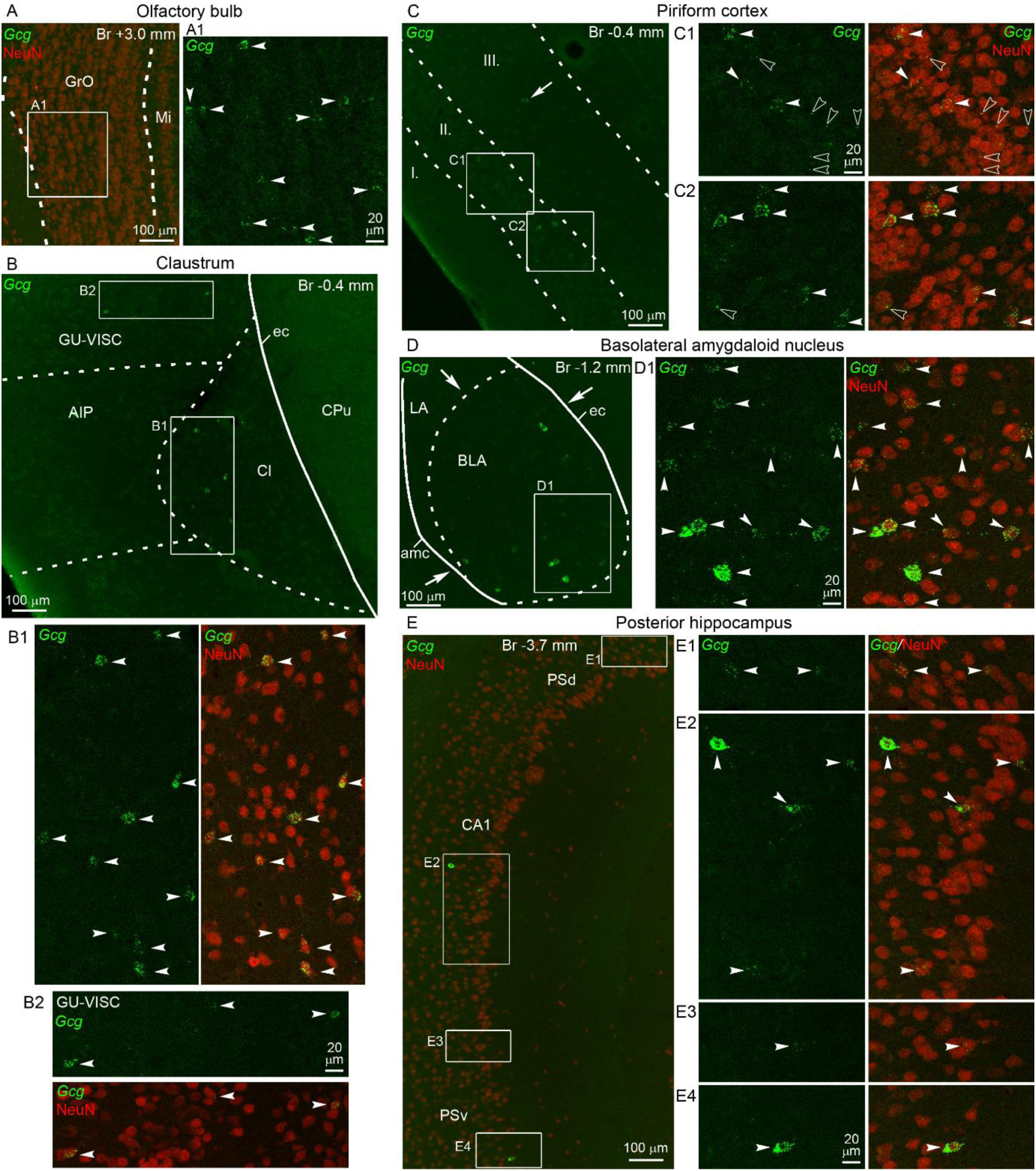
*Gcg*-expressing neuron populations in the cerebrum. Combination of *Gcg* FISH (green) with NeuN IF (neuronal marker, red). Low magnification images (*Gcg* or merged *Gcg*/NeuN signals to aid identification of brain regions) show *Gcg* mRNA-expressing neuron populations in the (**A**) olfactory bulb, (**B**) claustrum and adjacent cortex, (**C**) piriform cortex, (**D**) rostral part of the basolateral amygdaloid nucleus, and (**E**) posterior hippocampus of male C57BL/6J mice. Boxed areas are shown in higher magnification confocal images. Merged *Gcg*/NeuN images illustrate that the *Gcg* FISH signal is localized in neuronal cell bodies. Arrowheads indicate *Gcg*-expressing neurons. Note that *Gcg* mRNA levels vary among cells within a population. Open arrowheads in **C1** and **C2** point to piriform cortex neurons with 1 or 2 hybridization dots. The arrow in **C** points to a *Gcg* neuron in layer III of the piriform cortex. Arrows in **D** indicate the perimeter inside which *Gcg*-expressing neurons are located, which corresponds to the BLA border. Bregma levels are indicated. Abbreviations: I-III., layers I-III. of the piriform cortex; AIP, agranular insular cortex, posterior part; amc, amygdalar capsule; BLA, basolateral amygdaloid nucleus; CA1, hippocampal CA1 field; Cl, claustrum; CPu, caudate putamen; ec, external capsule; GrO, granule cell layer of the olfactory bulb; GU-VISC, gustatory-visceral cortex; LA, lateral amygdaloid nucleus; Mi, mitral cell layer of the olfactory bulb; PSd, dorsal prosubiculum; PSv, ventral prosubiculum.

**Figure 2.**
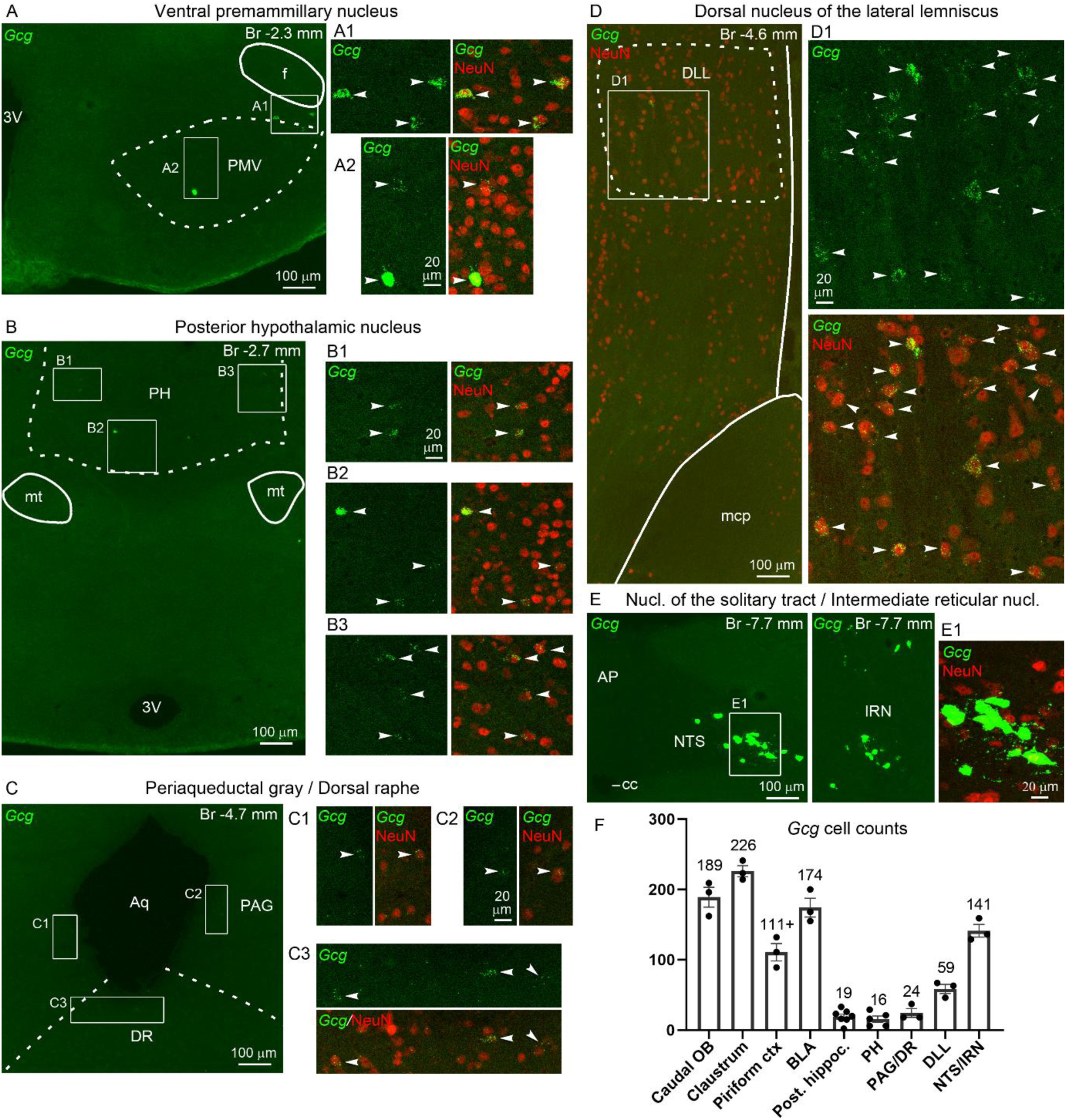
*Gcg*-expressing neuron populations in the hypothalamus and brainstem. Combination of *Gcg* FISH (green) with NeuN IF (neuronal marker, red). Low magnification images (*Gcg* or merged *Gcg*/NeuN signals to aid identification of brain regions) show *Gcg* mRNA-expressing neurons in the (**A**) ventral premammillary nucleus, (**B**) posterior hypothalamic nucleus, (**C**) periaqueductal gray and dorsal raphe, (**D**) dorsal nucleus of the lateral lemniscus, and (**E**) nucleus of the solitary tract and intermediate reticular nucleus in male C57BL/6J mice. Higher magnification confocal images of the boxed areas show *Gcg* and merged *Gcg*/NeuN signals, demonstrating the localization of *Gcg* mRNA in neuronal perikarya. Arrowheads indicate *Gcg*-expressing neurons. Bregma levels are indicated. (**F**) Cell counts of *Gcg*-expressing neurons in the different populations (counted in every 12th 16 µm thick section). In the OB, cells were counted in its caudal part, between Bregma +2.5 to 3.3 mm (4-5 sections). In the piriform cortex, the + sign indicates that this number is a substantial underestimate (see section 2*.1*). Cells were counted in three *ad libitum* fed male C57BL/6J mice (10, 15 and 20 wks). To increase sample size for two smaller *Gcg* neuron populations, we included cell numbers in the PH from two more mice (12 wks; not part of the fasting experiment), and in the posterior hippocampus from four additional mice (14-15 wks, fed controls from the fasting experiment). Mean values are shown above columns. Abbreviations: 3V, third ventricle; Aq, cerebral aqueduct; AP, area postrema; cc, central canal; DLL, dorsal nucleus of the lateral lemniscus; DR, dorsal raphe nucleus; f, fornix; IRN, intermediate reticular nucleus; mcp, middle cerebellar peduncle; mt, mammillothalamic tract; NTS, nucleus of the solitary tract; PAG, periaqueductal gray; PH, posterior hypothalamic nucleus; PMV, ventral premammillary nucleus.

#### 2.1.1 Cerebrum

##### OB

*Gcg*-expressing neurons were distributed in the granule cell layer of the OB and had uniformly low/moderate *Gcg* mRNA levels (**Fig 1A**, **A1**). We did not observe *Gcg* expression in the glomerular layer. Given the long rostro-caudal extent of the OB, this population contains the largest number of *Gcg*-expressing neurons in the brain. Our cell counts included here only the caudal OB, from Bregma +2.5 to 3.3 mm (4-5 sections; **Fig 2F**), due to tissue loss from the rostral/mid OB.

##### Claustrum

One of the largest *Gcg* neuron population by cell number (**Fig 2F**) was distributed in the claustrum and adjacent cortical areas (**Fig 1B-B2**). *Gcg* neurons were observed through the entire antero-posterior length of the claustrum (Bregma + 2.0 to -1.1 mm), in both the medial and lateral parts. Few cells were observed in the gustatory-visceral cortex, agranular insular cortex, or ventrally in the dorsal endopiriform nucleus. According to a narrower definition of the claustrum [32], more cells of this population might belong to layer 6 of the overlying cortical areas. Neurons of this population generally had low to moderate, but occasionally high *Gcg* mRNA levels.

##### Piriform cortex

*Gcg* neurons were observed predominantly in layer II (pyramidal layer), and occasionally in layer III of the piriform cortex (**Fig 1C-C2**), through its entire antero-posterior extent (Bregma +2.5 to -2.8 mm), but most commonly from Bregma +0.3 to -1.3 mm. *Gcg* mRNA levels were heterogeneous among cells, mostly low to moderate, but high-level expression also occurred. This is also a relatively large *Gcg* neuron population (**Fig 2F**), even though our cell counts substantially underestimate its real cell number, partly due to a stricter cell counting criterion applied here (see section *4.13*) and also because cells with 1-2 hybridization dots were more common in this population. It was evident in completely background-free sections that these single or double dots were always associated to neuronal cell bodies (**Fig 1C1**, **C2**).

##### BLA

A large population of *Gcg*-expressing neurons was observed specifically in the rostral part of the BLA (**Fig 1D**, **D1**), between Bregma -0.7 and -1.3 mm. *Gcg* mRNA levels were somewhat variable, generally moderate or low, but some cells expressed high levels.

##### Posterior hippocampus

A small population of *Gcg*-expressing neurons was observed in a narrow antero-posterior zone of the hippocampus, approximately from Bregma -3.5 to -4.1 mm (**Fig 1E-E4**). Within this zone, *Gcg* neurons were distributed in the CA1 field, and dorsal, caudal or occasionally ventral to it, in the prosubiculum, as defined in Ref. [33]. Hippocampal *Gcg* neurons appeared to be pyramidal cells, typically positioned close to the medial edge of the pyramidal layer. *Gcg* mRNA levels varied considerably among these cells, but moderate or low levels were more common than high-level expression.

#### 2.1.2 Hypothalamus

*Ventral premammillary nucleus (PMV).* Small and varying numbers of *Gcg* neurons were observed in the PMV and its vicinity in male C57BL/6J mice (**Fig 2A-A2**). We observed *Gcg* neurons in 14 out of the 18 male C57BL/6J mice examined in this study, including both fed and fasted animals. *Gcg* neuron numbers varied from 1 to 32 (counted in 2-3 sections containing the PMV in the series of 1 in 12 sections) but only five brains had >10 *Gcg* neurons (**Fig S1A**, **B-B5**), which we considered the minimum number for a cell population. *Gcg* expression was not observed in the PMV in any of the eight female C57BL/6J mice studied. Occasional *Gcg* neurons (1-3 cells) were observed in only 6 out of the 17 male FVB/Ant mice examined.

*Posterior hypothalamic nucleus (PH)*: A small *Gcg* neuron population was distributed in the posterior part of the PH, caudal of Bregma -2.3 mm (**Fig 2B-B3**). *Gcg* expression varied among cells, but most had low *Gcg* mRNA levels. This population is described in more detail in section *2.2*.

#### 2.1.3 Midbrain and hindbrain

##### Periaqueductal gray (PAG) and dorsal raphe (DR)

A small population of *Gcg* neurons was distributed ventral/ventrolateral to the cerebral aqueduct, between Bregma -3.5 to -4.9 mm. The majority of these cells concentrated caudally, between Bregma -4.2 to -4.9 mm, in the ventrolateral PAG, and dorsal and ventral parts of the DR (**Fig 2C-C3**). Very low *Gcg* mRNA levels were characteristic of this population, although cells with moderate levels also occurred.

##### Dorsal nucleus of the lateral lemniscus (DLL)

A medium-sized neuron population expressing generally low to moderate *Gcg* mRNA levels was observed in the DLL (**Fig 2D**, **D1**).

##### NTS/IRN

Medullary *Gcg* neurons were distributed in the NTS and IRN (**Fig 2E**, **E1**), with few cells in the raphe obscurus, as previously described [6; 26]. Almost all of these cells expressed very high *Gcg* mRNA levels, only occasional cells expressed moderate levels.

#### 2.1.4 Scattered Gcg neurons in other areas

We observed scattered *Gcg* neurons consistently or often in several other brain regions. These include the anterior olfactory nucleus, lateral/ventral orbital cortex, somatosensory cortex, dorsal and ventral endopiriform nuclei, collectively in various amygdala nuclei (cortex-amygdala transition zone, anterior cortical amygdaloid nucleus, basomedial amygdaloid nucleus, posterior and ventral parts of the BLA, medial amygdaloid nucleus, amygdalopiriform transition area, posterolateral cortical amygdaloid nucleus), zona incerta, collectively in mid-caudal hypothalamic nuclei (dorsomedial, ventromedial or arcuate nucleus), and the parabrachial nucleus. *Gcg* neurons were occasionally observed in the primary motor cortex, lateral entorhinal cortex, dentate gyrus, ventral pallidum, rostral perifornical area, parasubthalamic nucleus, red nucleus, inferior colliculus and medial vestibular nucleus.

### 2.2 Fasting markedly increases Gcg mRNA expression in the PH

Based on a report that glicentin-immunoreactive cells were detected in the hypothalamic arcuate and ventromedial nuclei of fasted rats [25], we studied the effect of 30h fasting on *Gcg* expression. We first compared brain-wide *Gcg* expression in fasted male and female C57BL/6J, and male FVB/Ant mice (n=3 from each) to control mice. Fasting visibly increased *Gcg* expression in the PH, which we examined further. No obvious change was observed in the other populations, although in NTS/IRN neurons the FISH signal was too high to accurately assess any alterations. We did not observe fasting-induced *Gcg* expression in the ventromedial and arcuate nuclei.

#### 2.2.1 The PH Gcg neuron cluster continues in the midbrain

Increased *Gcg* expression in the PH of fasted mice revealed that these cells constitute a medium-sized population that extends into the midbrain (**Fig 3**). Aside from few cells in the rostral PH, the bulk of this population was distributed in the caudal PH, from Bregma -2.3 to -3.0 mm. In its most caudal part, *Gcg* neurons were distributed more broadly than what is shown as PH in reference atlases [34; 35], and occasionally observed in the supramammillary nucleus. *Gcg* neurons continued caudally in smaller number, within and around the midbrain interfascicular nucleus (IFN) that sits atop the interpeduncular nucleus (**Fig 3B**, **E**). Since fasting increased *Gcg* mRNA expression in both the PH and IFN, we counted these cells collectively as one population. The proportion of IFN *Gcg* neurons was on average 11% of all *Gcg* neurons in the combined PH/IFN. Few *Gcg* neurons could be observed even more caudally in the midline linear raphe nuclei. These were not included in the analysis, but were conspicuous in fasted FVB/Ant mice, and also observed in some fed FVB/Ant, and fasted C57BL/6J mice.

**Figure 3.**
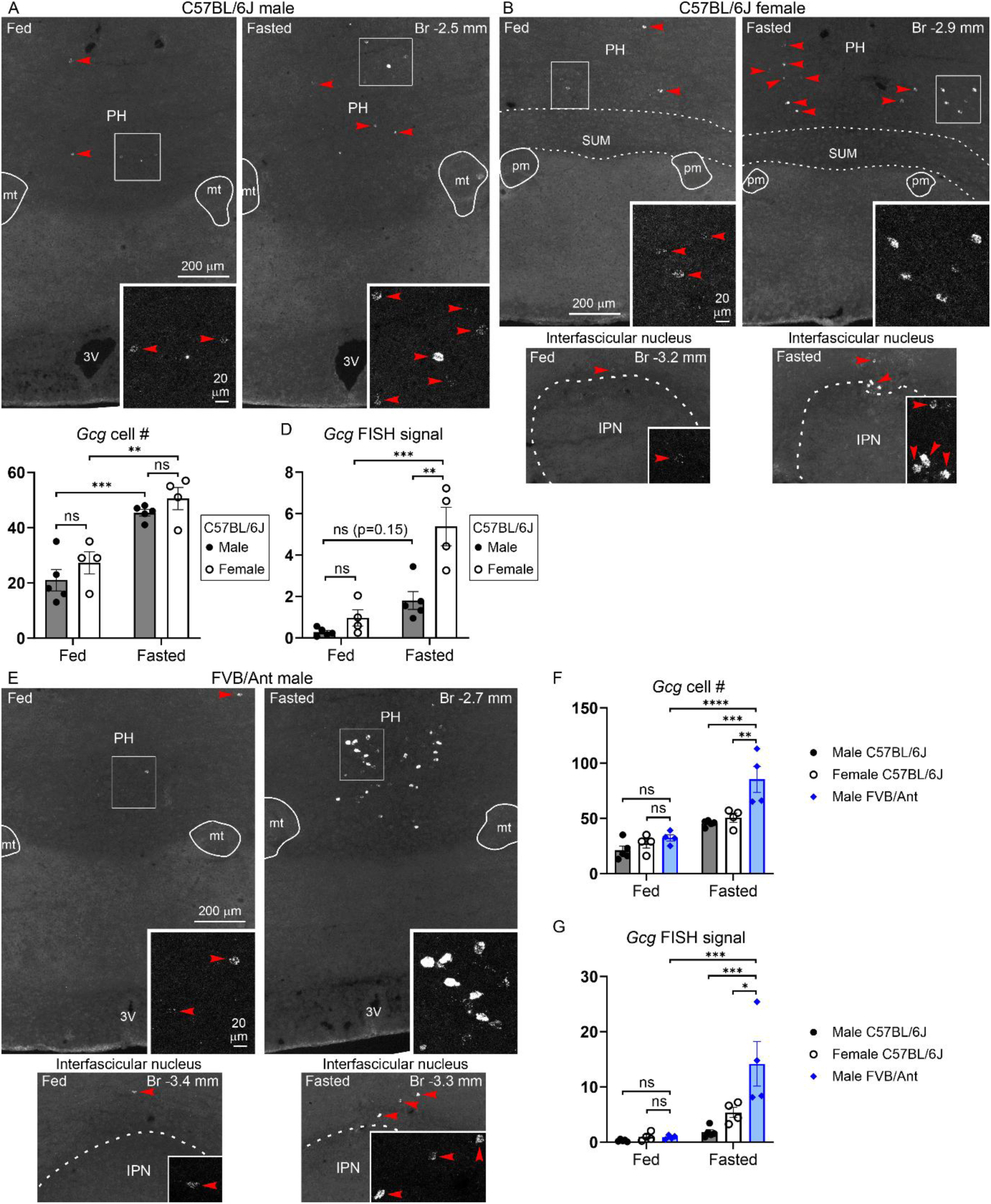
Fasting increases *Gcg* expression in the PH/IFN. (**A**, **B**, **E**) FISH shows the effect of 30h fasting on *Gcg* mRNA expression in the PH of male C57BL/6J mice (**A**), the PH and IFN of female C57BL/6J mice (**B**), and the PH and IFN of male FVB/Ant mice (**E**). Bregma levels are indicated. Red arrowheads indicate *Gcg* neurons; the boxed areas and IFN *Gcg* neurons indicated by arrowheads are shown at higher magnification in the insets. (**C, D**, **F**, **G**) Graphs showing the number of detected *Gcg* neurons (**C, F**), and *Gcg* FISH signal measured by image analysis (integrated density in arbitrary units) (**D, G**) in the combined PH/IFN (counted/measured in every 12th 16 µm thick section). Group sizes: male C57BL/6J: n=5 per group (14-15 wks); female C57BL/6J: n=4 per group (10-11 wks); male FVB/Ant: n=4 per group (16 wks). Two-way ANOVA and Tukey’s multiple comparison’s test were used to compare data between male and female C57BL/6J mice (**C**, **D**). One-way ANOVA and Tukey’s multiple comparison’s test were used to compare data between male FVB/Ant and male and female C57BL/6J mice (**F**, **G**). P values of the Tukey’s tests between individual groups are indicated: ns: non-significant, *<0.05, **<0.01, ***<0.001, ****<0.0001. Abbreviations: 3V, third ventricle; IPN, interpeduncular nucleus; mt, mammillothalamic tract; PH, posterior hypothalamic nucleus; pm, principal mammillary tract; SUM, supramammillary nucleus.

#### 2.2.2 Sex and strain differences in fasting-induced Gcg expression in the PH/IFN

Fasting markedly increased *Gcg* mRNA expression in the PH/IFN in all groups of mice, but more strikingly in FVB/Ant, than in C57BL/6J mice (**Fig 3**; n=4-5 per experimental group). In *ad libitum* fed C57BL/6J mice, *Gcg* expression was more variable and somewhat higher in females than in males, but not significantly different (**Fig 3A-D**). Fasting significantly increased the number of detected *Gcg* neurons to approximately 2-fold [two-way ANOVA, main effect: fasting, F(1, 14) = 49.40, p<0.0001], with no statistical difference between the sexes [two-way ANOVA, main effect: sex, F(1, 14) = 2.803, p=0.1163] (**Fig 3C**). However, the fasting-induced increase in *Gcg* FISH signal, a proxy for total *Gcg* mRNA levels, showed an interaction with sex [two-way ANOVA, main effect: fasting, F(1, 14) = 34.45, p<0.0001; main effect: sex, F(1, 14) = 17.65, p=0.0009; interaction, F(1, 14) = 8.091, p=0.0130) (**Fig 3D**). This increase was approximately 6-fold in both sexes, although statistically significant only in females (females: p=0.0002; males: p=0.1545, Tukey’s multiple comparisons test) (**Fig 3D**). Fasted females had significantly, and noticeably, higher *Gcg* FISH signal than fasted males (p=0.0010, Tukey’s test) (**Fig 3A**, **B**, **D**).

In *ad libitum* fed male FVB/Ant mice, *Gcg* mRNA expression was comparable to, and not significantly different from that of C57BL/6J mice (Tukey’s multiple comparisons following significant one-way ANOVA, p<0.0001, for both *Gcg* cell # and FISH signal) (**Fig 3E-G)**. In male FVB/Ant mice, fasting significantly increased the number of detected *Gcg* neurons to almost 3-fold (p<0.0001, Tukey’s), and the *Gcg* FISH signal to 16-fold (p<0.0002, Tukey’s) (**Fig 3E-G**). The values in both measures were significantly higher than in fasted C57BL/6J mice, either male (*Gcg* cell #: p=0.0004; FISH signal: p=0.0002) or female (*Gcg* cell #: p=0.0033; FISH signal p=0.0119) (**Fig 3F**, **G**). In several cells, *Gcg* mRNA reached such levels that the FISH signal filled the cytoplasm, similarly to medullary *Gcg* neurons (**Fig 3E**). We confirmed in another, 48h fasting experiment, that such high level of *Gcg* expression in the PH is typical in fasted FVB/Ant mice (**Fig S2**).

### 2.3 PG and GLP-1 synthesis in Gcg-expressing neuron populations

We used IF to examine whether PG and GLP-1 are synthesized in *Gcg*-expressing neuron populations. To detect PG, we used a rabbitized monoclonal antibody that binds to a mid-molecule GLP-1 epitope, also present in GLP-1 precursors (PG antibody) (**Fig S3**). For GLP-1, we used two mouse monoclonal antibodies: one that is specific to the amidated C-terminus present in both the biologically active GLP-1(7-36)amide and the inactive precursor GLP-1(1-36)amide (GLP-1^C^ antibody); and another that binds the free N-terminus of the bioactive forms, GLP-1 (7-36)amide and GLP-1 (7-37) (GLP-1^N^ antibody). We applied a modified IF protocol that significantly enhanced PG and GLP-1 detection (see *Methods* and **Fig S4**); PG IF with the standard protocol is shown in **Fig S5**.

In the medulla, the PG antibody intensely labeled both the cell bodies of *Gcg* neurons in the NTS/IRN, and axons (**Fig 4A**). The GLP-1^C^ antibody only faintly labeled these perikarya but intensely labeled axons (**Fig 4A**). GLP-1^N^ IF also resulted in faint perikaryal, but intense axonal labeling (**Fig S3E**, **F**). This suggests that the PG antibody signal in cell bodies predominantly represents GLP-1 precursors, such as PG or major proglucagon fragment. Conversely, GLP-1 is present primarily in axons, and less concentrated in cell bodies.

**Figure 4.**
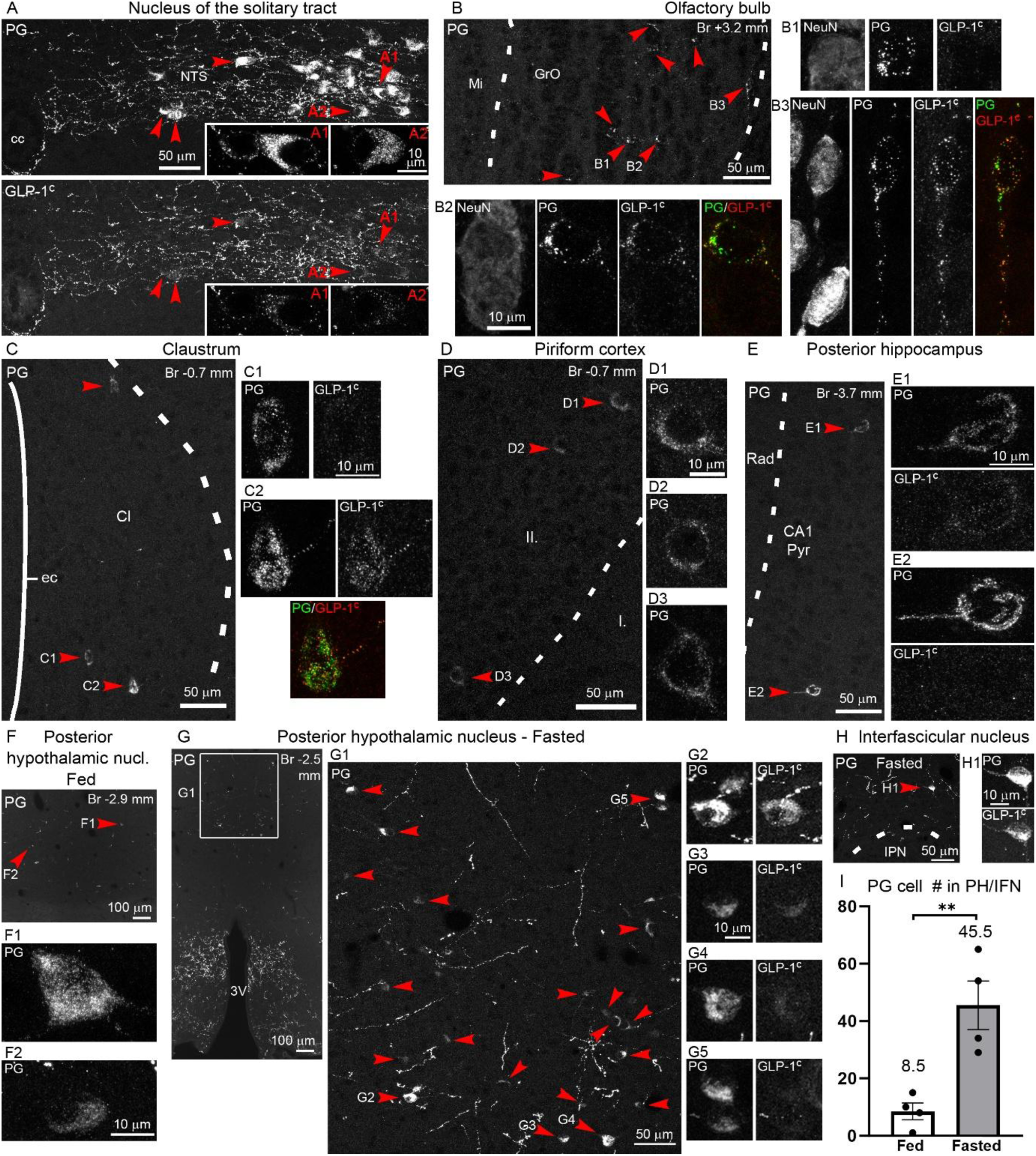
PG and GLP-1 synthesis in *Gcg*-expressing neuron populations. Triple-label IF for PG/GLP-1^C^/NeuN in male FVB/Ant mice (11-12 wks). Arrowheads indicate PG neurons. (**A**) PG and GLP-1^C^ signals (confocal Z-projections) in the NTS, shown in separate panels: cell bodies display intense PG, and faint GLP-1^C^ signals (arrowheads indicate examples). Insets show single optical sections of the numbered cells. (**B**) PG neurons in the OB. High magnification images show the NeuN, PG and GLP-1^C^ signals of the numbered cells: cell **B1** is GLP-1^C^-negative, cells **B2** and **B3** are GLP-1^C^-positive (merged PG/GLP-1^C^ panels included). (**C**) PG neurons in the claustrum. High magnification images show the PG and GLP-1^C^ signals of the numbered cells: cell **C1** is GLP-1^C^-negative, cell **C2** is GLP-1^C^-positive (merged PG/GLP-1^C^ panel included). (**D**) PG neurons in the piriform cortex. GLP-1^C^ signals are not shown, as these cells are GLP-1^C^-negative. (**E**) PG neurons in the posterior hippocampus. High magnification images show their PG and GLP-1^C^ signals: cell **E1** is light positive for GLP-1^C^, cell **E2** is GLP-1^C^-negative. (**F**) PG neurons in the PH of a fed mouse, shown also in high magnification images. (**G**) Low magnification image from a fasted mouse shows the location of PG neurons (boxed area) in the PH. Higher magnification confocal image of this area (**G1**) shows several PG neurons, with varying labeling intensities. Magnified images of the numbered cells show their PG and GLP-1^C^ signals in separate panels: **G2:** strong GLP-1^C^-positive cell next to a GLP-1^C^-negative cell; **G3:** light GLP-1^C^-positive cell; **G4**: minimally GLP-1^C^-positive cell; **G5**: two GLP-1^C^-negative cells. (**H)** A PG neuron in the IFN of a fasted mouse. High magnification images show that it is intensely positive for both PG and GLP-1^C^. (**I**) PG neuron numbers (counted in every 4th 25 µm thick section) in the PH/IFN of fed and 30h fasted male FVB/Ant mice (n=4 each, 12 wks). Mean values are shown above columns. Statistical significance: **<0.01, t-test. Abbreviations: I-II, layers of the piriform cortex; 3V, third ventricle; CA1 Pyr, CA1 pyramidal layer; cc, central canal; Cl, claustrum; ec, external capsule; GrO, granule cell layer of the olfactory bulb; IPN, interpeduncular nucleus; Mi, mitral cell layer of the olfactory bulb; NTS, nucleus of the solitary tract; Rad, radiatum layer of the hippocampus.

We examined the rest of the brain in male FVB/Ant mice (11-15 wks) with triple-labeling for PG/GLP-1^C^/NeuN, the latter to confirm that the PG labeling is present in neuronal perikarya. We detected PG-containing neurons in each area where *Gcg* mRNA is expressed at low levels, but fewer than what was detected by FISH. Most of these PG neurons were lightly or moderately labeled. **Fig S6E** shows the PG neuron counts through serial sections per population; here we will state the cell numbers observed per section. PG neuron number and distribution were similar in C57BL/6J mice, with noted exceptions. GLP-1^N^ immunolabeling were examined only in the OB and PH/IFN, with triple-labeling for PG/GLP-1^N^/NeuN.

#### 2.3.1 Cerebrum

The granule cell layer of the OB contained the highest number of PG neurons (∼20-50 per section per side). These perikarya contained PG in distinct granules/vesicles (**Fig 4B-B3**), mirroring the Golgi apparatus morphology of OB granule cells [36]. The PG-containing vesicles also extended into neurites. In numerous cells, the GLP-1^C^ antibody co-labeled the PG-containing granules/vesicles (**Fig 4B2**, **B3**), but GLP-1^C^-negative cells were also observed (**Fig 4B1**). We did not detect GLP-1^N^ signal in OB PG neurons.

The claustrum contained PG neurons through its rostrocaudal extent: 1-7 cells per section per side were observed in nearly every section (**Fig 4C-C2**). Few PG neurons were observed in the gustatory-visceral cortex. GLP-1^C^ was detected in a minority of claustrum PG neurons (7.9 ± 2.0%, n=3) (**Fig 4C2**), often just above background level. PG neurons were observed in layer II of the piriform cortex (rarely in layer III), particularly between Bregma +0.6 and -1.3 mm, with 1-7 PG neurons per section per side detected in most sections (**Fig 4D-D3**). Piriform cortex PG neurons were GLP-1^C^-negative. In the posterior hippocampus, up to 3 PG neurons per section per side were observed (**Fig 4E-E2**). The PG signal formed the typical Golgi apparatus shape of hippocampal pyramidal neurons [37; 38]. Some of these cells had intense PG signal, and few were faint positive for GLP-1^C^ (**Fig 4E1**). In the BLA, very few PG neurons were detected relative to the number of *Gcg* mRNA-expressing neurons in this nucleus: up to 4 per section per side, some GLP-1^C^-positive (**Fig S6A-A3**). In the BLA of female C57BL/6J mice, however, we detected significantly more PG neurons, up to 20 per section per side (**Fig S6B**). Of note, PG signal was localized mainly to the Golgi apparatus in the vast majority of claustrum, piriform cortex, BLA and hippocampus neurons (**Figs 4C1**, **D1-3**, **E1**, **E2**, **S6A1**, **A2**, **B1**, **B2**). More dispersed PG signal, suggesting localization also in secretory vesicles, was observed in few cells (**Figs 4C2**, **S6A3**). PG neurons were occasionally observed in the somatosensory cortex, lateral orbital cortex, and dorsal endopiriform nucleus.

#### 2.3.2 Hypothalamus and brainstem

In the PH/IFN of *ad libitum* fed mice, a small number of lightly or moderately labeled PG neurons were detected (**Figs 4F-F2**, **I**, **S6E**). Significantly more PG neurons, including several intensely labeled, were observed in 30h fasted mice (p=0.0061, t-test) (**Fig 4G-I**). Occasional PG neurons were observed in the linear raphe nuclei. Approximately 10-20% of PH/IFN PG neurons were GLP-1^C^-positive (fed: 11.5 ± 4.7%, fasted: 14.0 ± 2.5%, n=4 each), generally the cells with more intense PG labeling (**Fig 4G2-H1**). We occasionally detected light GLP-1^N^ signal in the most intensely labeled PG neurons. A replicate of the fasting experiment with similar upregulation of PG synthesis is shown in **Fig S5I-K**.

In the PMV, only a single PG neuron was found among the 12 male FVB/Ant brains examined. In male C57BL/6J mice (n=3), the number of PG neurons in the PMV varied among brains from none to several cells, confirming the FISH results (**Fig S1C-E1**). The more intensely labeled PG neurons were GLP-1^C^-positive (**Fig S1D-E1**).

In the PAG/DR, only single or no PG neurons were observed through serial sections, except a slightly older brain (15 wks), in which we observed several PG neurons (**Fig S6C-C4**, **E**). In the DLL, up to 7 PG neurons per section per side were observed, most with light labeling (**Fig S6D-E**). PG neurons were occasionally GLP-1^C^-positive in the DR and DLL (**Fig S6C2**, **D1**). In the hypothalamic and brainstem populations, a larger portion of PG neurons had PG signal dispersed in the cell body and even dendrites, suggesting localization in secretory vesicles (**Figs 4F1**, **G1-H1**, **S1D2**, **E1**, **S6C1**, **C2**, **D1**).

### 2.4 Characterization of Gcg-Cre mice

To understand whether BAC *Gcg*-Cre mice [27] can be used to study the newly identified *Gcg* neuron populations, we characterized transgene expression in *Gcg*-Cre;tdTomato mice, generated by crossing *Gcg*-Cre with tdTomato-expressing Ai9 Cre reporter mice. The initial description of these mice mentioned few tdTomato-positive neurons in various brain regions, including the piriform cortex, PH, PMV, and PAG/DR [27]. In addition to double heterozygotes (*Gcg*-*Cre*^/+^;*tdTom*^/*+*^), we also generated double homozygote mice (*Gcg*-*Cre*^/*Cre*^;*tdTom*^/*tdTom*^) to increase the number of tdTomato-labeled neurons.

#### 2.4.1 Distribution of tdTomato cells in Gcg-Cre;tdTomato mice

Significant populations of tdTomato neurons were observed in the NTS/IRN, granule cell layer of the OB, PH/IFN, and PAG/DR, in both genotypes (**Fig 5A-E**). In the DLL, only *Gcg*-*Cre*^/*Cre*^;*tdTom*^/*tdTom*^ mice had a modest but significant number of tdTomato neurons. Rare or few tdTomato neurons were observed in the piriform cortex (mostly layer III), posterior hippocampus and PMV, and almost none in the claustrum and rostral BLA. In Supplementary Table 1 we summarized tdTomato cell counts from 13 *Gcg-*Cre;tdTomato mice of different genotype, sex, age and feeding status. All of these variables appear to influence the number of tdTomato neurons in certain populations. *Gcg*-*Cre*^/*Cre*^;*tdTom*^/*tdTom*^ mice had more tdTomato neurons than *Gcg*-*Cre*^/+^;*tdTom*^/*+*^ mice in all regions. Among *Gcg*-*Cre*^/+^;*tdTom*^/*+*^ mice, females had more tdTomato neurons in the PH/IFN than males. The number of tdTomato neurons tends to increase with age in the PAG/DR and following 24h fasting in the PH/IFN.

**Figure 5.**
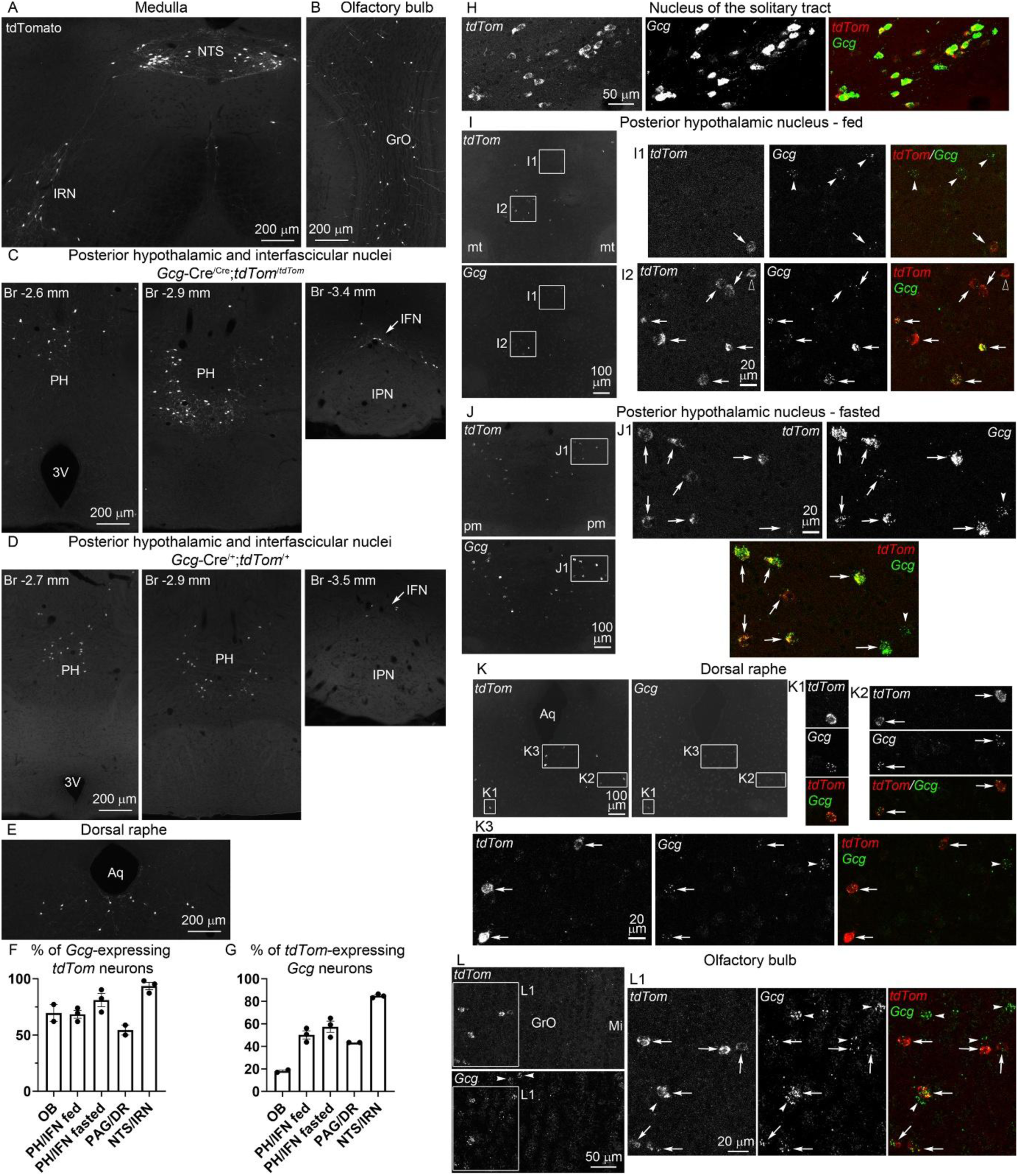
Characterization of tdTomato expression in *Gcg*-Cre;tdTomato mice. (**A-E**) Distribution of tdTomato neurons (native fluorescence) in the (**A**) NTS/IRN, (**B**) granule cell layer of the OB, (**C-D**) PH/IFN (Bregma levels indicated), and (**E**) dorsal raphe. **A-C**, **E** are from *Gcg*-*Cre*^/*Cre*^;*tdTom*^/*tdTom*^ mice, **D** is from a *Gcg*-*Cre*^/*+*^;*tdTom*^/*+*^ mouse. (**F**) Percentage of *Gcg*-expressing *tdTom* neurons in the dual FISH studies (n=2 or 3). (**G**) Percentage of *tdTom*-expressing *Gcg* neurons in the dual FISH studies (n=2 or 3). (**H-L**) Dual FISH for *tdTom* and *Gcg* mRNAs. The low magnification images show *tdTom* and *Gcg* signals in separate panels; the boxed areas are shown in higher magnification confocal images, including merged *tdTom* (red)/*Gcg* (green) panels. (**H**) Confocal images of the NTS, where co-expression of *tdTom* and *Gcg* mRNAs is virtually complete. (**I**) In the PH of fed mice, the majority of *tdTom* neurons express *Gcg* mRNA (arrows), but often at low levels. *TdTom* neurons with no *Gcg* hybridization signal (open arrowhead in **I2**), or *Gcg* neurons with no *tdTom* mRNA (arrowheads in **I1**) also occur. (**J**) In the PH of fasted mice, *tdTom* neurons express higher *Gcg* mRNA levels. (**K**) In the dorsal raphe, examples of dual-labeled *tdTom*/*Gcg* neurons (arrows), and a *Gcg* neuron without *tdTom* mRNA (arrowhead) are shown. (**L**) In the OB, most *tdTom* neurons express *Gcg* mRNA (arrows), but the majority of *Gcg* neurons do not express *tdTom* mRNA (arrowheads). Abbreviations: 3V, third ventricle; Aq, aqueduct; GrO, granule cell layer of the olfactory bulb; IFN, interfascicular nucleus; IPN, interpeduncular nucleus; IRN, intermediate reticular nucleus; Mi, mitral cell layer of the olfactory bulb; mt, mammillothalamic tract; NTS, nucleus of the solitary tract; PH, posterior hypothalamic nucleus; pm, principal mammillary tract.

We observed two other sizable populations of tdTomato neurons: one distributed in the broad amygdala region, the other in the zona incerta (Supplementary Table 1). Few, scattered tdTomato neurons were observed in other brain regions in *Gcg*-*Cre*^/+^;*tdTom*^/*+*^ mice. The distribution of these cells, including the amygdala region and zona incerta, largely corresponded to the pattern of scattered *Gcg* mRNA-expressing neurons detected with FISH. Scattered tdTomato neurons were observed in many other brain regions in *Gcg*-*Cre*^/*Cre*^;*tdTom*^/*tdTom*^ mice. Of note, in some brains we observed few tdTomato-positive non-neuronal cells, including hypothalamic tanycytes, and glial cells in the caudal hypothalamus, PAG and medulla. We observed two cases what appeared to be tdTomato activation in neural progenitor cells and subsequent neurogenesis: in the hypothalamic premammillary/mammillary nuclei, and in the medullary spinal trigeminal nucleus, reminiscent of the clonal expansions reported in *Gcg^iCre^* mice [39].

#### 2.4.2 TdTomato labels four Gcg-expressing neuron populations

To verify tdTomato expression in *Gcg* neurons, we used dual FISH, or additionally dual IF. Cells in the FISH experiments were counted on every 12th 16 µm thick section through a neuron population; co-expression percentages are summarized in **Fig 5F**, **G**. In the NTS/IRN, 93.6 ± 3.3% of *tdTom* neurons expressed *Gcg* (of 121.3 ± 5.2 *tdTom* neurons, from 3 female *Gcg*-*Cre*^/*+*^;*tdTom*^/*+*^ mice, 13-14 wks) (**Fig 5H**). Conversely, 84.8 ± 1.0% of *Gcg* neurons expressed *tdTom* (of 133.7 ± 5.4 *Gcg* neurons). Of note, co-expression was almost complete in the NTS, as single-labeled *Gcg* or *tdTom* neurons were observed primarily in the IRN, in agreement with the initial characterization [27].

Dual FISH in the PH/IFN was performed in both *ad libitum* fed and 30h fasted *Gcg*-*Cre*^/*+*^;*tdTom*^/*+*^ female mice (n=3 each, 13-14 wks) (**Fig 5I**, **J**). In fed females, 68.5 ± 3.5% of *tdTom* neurons expressed *Gcg* (of 54.0 ± 4.0 *tdTom* neurons), while 81.0 ± 5.8% expressed *Gcg* in fasted females (of 59.0 ± 6.6 *tdTom* neurons). Conversely, 50.1 ± 3.6% of *Gcg* neurons expressed *tdTom* in fed females (of 74.0 ± 5.7 *Gcg* neurons), and 57.4 ± 5.0% expressed *tdTom* in fasted females (of 83.3 ± 8.6 *Gcg* neurons). Here we considered *tdTom* neurons even with a single *Gcg* hybridization dot as *Gcg* positive; such cells accounted for 17.2 ± 1.8% of *tdTom/Gcg* neurons in fed, and 8.1 ± 2.1% in fasted mice. To study co-expression at the protein level, we performed triple IF for PG/tdTomato/NeuN in 30h fasted female *Gcg*-*Cre*^/*+*^;*tdTom*^/*+*^ mice (n=3, 10 wks; **Fig S7A-A2**). PG was expressed in 33.9 ± 9.2% of tdTomato neurons (of 102.7 ± 8.3 tdTomato neurons, counted on every 4th 25 µm thick section through the PH/IFN). Conversely, 72.4 ± 6.2% of PG neurons contained tdTomato (of 47.0 ± 12.1 PG neurons).

To assess *tdTom/Gcg* co-expression in the PAG/DR and granule cell layer of the OB, we examined a female (9 wks) and a male (13 wks) *Gcg*-*Cre*^/*Cre*^;*tdTom*^/*tdTom*^ mouse, as this genotype has markedly more tdTomato neurons in these nuclei than *Gcg*-*Cre*^/*+*^;*tdTom*^/*+*^ mice. In the PAG/DR, 50.0% and 58.8% of of *tdTom* neurons expressed *Gcg* (of 20 and 34 *tdTom* neurons, respectively), and in both mice 43.5% of *Gcg* neurons expressed *tdTom* (of 23 and 46 *Gcg* neurons) (**Fig 5K**). In the OB, 77.2% and 62.1% of *tdTom* neurons expressed *Gcg* (of 259 and 227 *tdTom* neurons) (**Fig 5L**). However, only 19.1% and 16.9% of *Gcg* neurons expressed *tdTom* (of 1046 and 835 *Gcg* neurons, counted on 10 sections from Bregma +2.7 to +4.6 mm). Here we also considered *tdTom* neurons with a single *Gcg* hybridization dot as *Gcg* positive: such cells accounted for 16.0% and 24.8% of *tdTom/Gcg* neurons. With IF we examined PG/tdTomato co-expression in the OB in two *Gcg*-*Cre*^/*Cre*^;*tdTom*^/*tdTom*^ female mice (8 wks) (**Fig S7B-B4**). PG was detected in 46.1 ± 0.9% of tdTomato neurons (159 ± 24 tdTomato neurons, counted in 10-15 unilateral sections). Conversely, most OB PG neurons lacked tdTomato, we estimate that <25% expressed tdTomato.

The *tdTom* neurons in the DLL of *Gcg*-*Cre*^/*Cre*^;*tdTom*^/*tdTom*^ mice (7 and 12 cells) did not contain *Gcg* mRNA.

### 2.5 Gcg-expressing neuron populations express PG-processing enzymes

We studied whether *Gcg* neuron populations express the main proteolytic enzymes that process PG: prohormone convertase 1/3 (PC1/3, encoded by the *Pcsk1* gene) and prohormone convertase 2 (PC2, encoded by *Pcsk2*). These enzymes process PG differently: PC1/3-mediated cleavage generates GLP-1 and GLP-2, while PC2 generates glucagon [40]. Because of the much higher concentration of GLP-1 than glucagon in the rat brain [41–43], PC1/3 has been thought to process PG in medullary GLP-1 neurons [44].

We applied dual FISH to detect *Pcsk1* or *Pcsk2* mRNA in *tdTom* neurons of *Gcg*-Cre;tdTomato mice. For each neuron population we used the same brains we examined for *Gcg*/*tdTom* co-expression. *Pcsk1* and *Pcsk2* expression patterns were very similar to the patterns described in rats [45]. In the NTS/IRN, nearly all *tdTom* neurons expressed *Pcsk1* (96.6 ± 1.2% of 111.7 ± 9.4 *tdTom* neurons; n=3), generally at moderate levels (**Fig 6A**, **A1**), though some cells had only 1-2 *Pcsk1* hybridization dots. The vast majority of *tdTom* neurons also expressed *Pcsk2* mRNA (79.8 ± 2.8% of 110.3 ± 11.5 *tdTom* neurons; n=3) (**Fig 6B**, **B1**). *Pcsk2* was expressed at lower levels, however, more often represented by 1-2 hybridization dots per cell.

**Figure 6.**
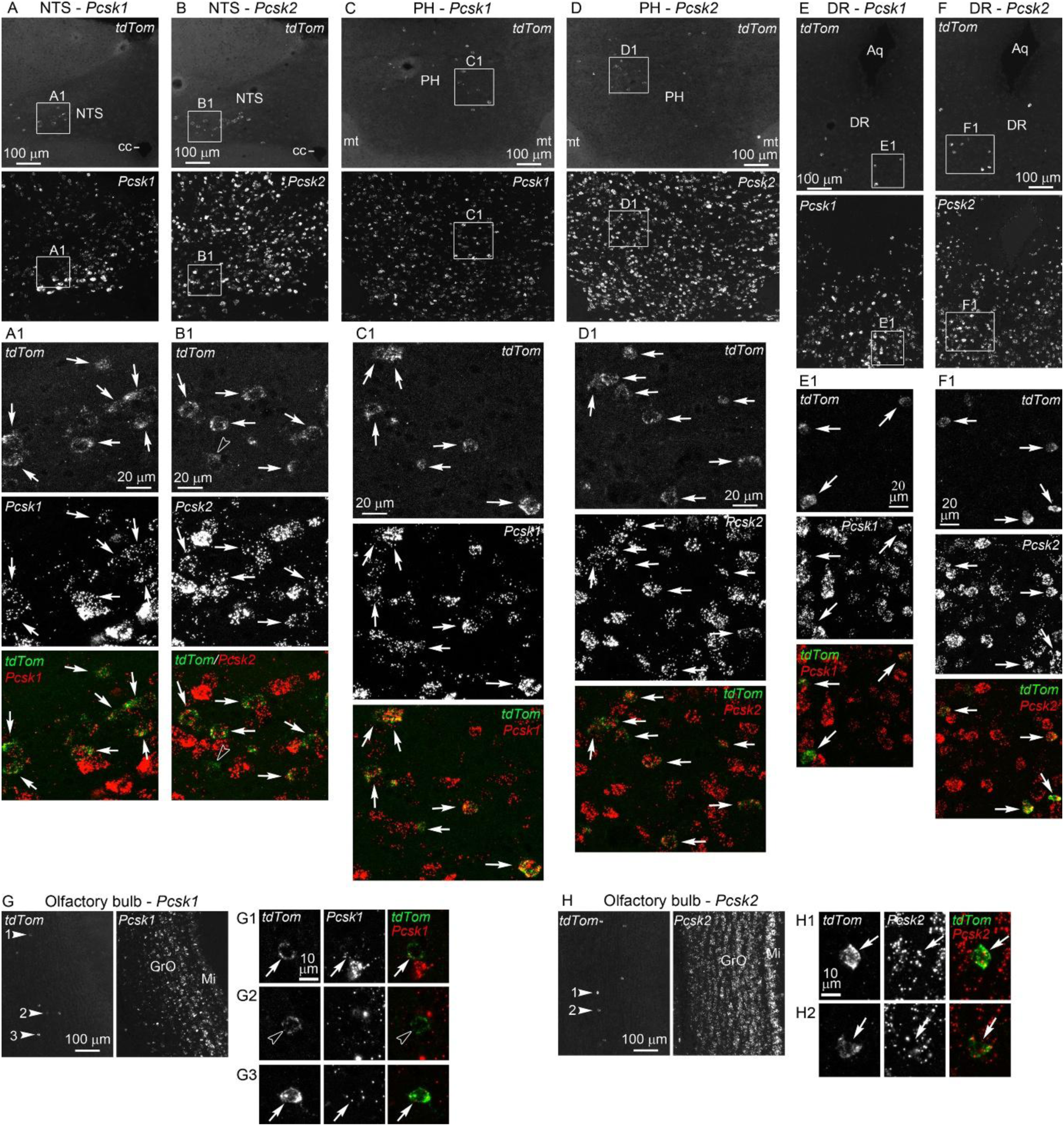
*Pcsk1* and *Pcsk2* expression in *tdTom*-labeled *Gcg*-expressing neuron populations. Dual FISH for *Pcsk1*/*tdTom* or *Pcsk2/tdTom* in the nucleus of the solitary tract (**A**, **B**), posterior hypothalamic nucleus (**C**, **D**), dorsal raphe (**E**, **F**) and olfactory bulb (**G**, **H**) of *Gcg*-Cre;tdTomato mice. Low magnification images show the individual FISH signals in separate panels. The boxed areas, or the *tdTom* neurons indicated by arrowheads in **G** and **H**, are shown in high magnification confocal images. The merged images show *tdTom* in green, *Pcsk1* or *Pcsk2* in red. Arrows indicate dual-labeled *Pcsk1*/*tdTom* or *Pcsk2/tdTom* neurons. Open arrowheads point to a *Pcsk2*-negative *tdTom* neuron in the NTS (**B**), and to a *Pcsk1*-negative *tdTom* neuron in the OB (**G2**). Abbreviations: Aq, aqueduct; DR, dorsal raphe; GrO, granule cell layer of the olfactory bulb; Mi, mitral cell layer of the olfactory bulb; mt, mammillothalamic tract; PH, posterior hypothalamic nucleus, NTS, nucleus of the solitary tract.

In the PH/IFN, virtually all *tdTom* neurons contained *Pcsk1* and *Pcsk2* mRNAs (**Fig 6C-D1**): 99.5 ± 0.5% expressed *Pcsk1* (of 53.7 ± 11.7 *tdTom* neurons, n=3), and 99.3 ± 0.7% expressed *Pcsk2* (of 51.3 ± 9.2 *tdTom* neurons, n=3). *Pcsk1* mRNA was expressed at moderate to high levels, *Pcsk2* mRNA at generally high levels. Fasting did not affect *Pcsk1* or *Pcsk2* expression in PH/IFN or NTS/IRN *tdTom* neurons (data not shown).

In the PAG/DR, all *tdTom* neurons expressed *Pcsk1* and *Pcsk2* mRNAs (of 28.5 ± 13.5 and 33.5 ± 12.5 *tdTom* neurons, respectively; n=2), both at generally high levels (**Fig 6E-F1**).

In the OB, 80.6 ± 1.0% of *tdTom* neurons expressed *Pcsk1* (of 233 ± 32 *tdTom* neurons; n=2). However, *Pcsk1* mRNA levels were generally low (**Fig 6G-G3**), with ∼20% of *Pcsk1*-positive *tdTom* neurons displaying only a single *Pcsk1* hybridization dot. All *tdTom* neurons in the OB expressed *Pcsk2,* at moderate levels (of 225.5 ± 0.5 *tdTom* neurons; n=2) (**Fig 6H-H2**).

To assess *Pcsk1* and *Pcsk2* expression in other *Gcg*-expressing populations not labeled by tdTomato in *Gcg*-Cre;tdTomato mice, we combined FISH with NeuN IF in male C57BL/6J mice (**Fig S8**). Virtually all claustrum neurons and piriform cortex pyramidal neurons expressed *Pcsk1* mRNA at moderate levels, and all BLA neurons at high levels. The vast majority of posterior hippocampus pyramidal cells expressed *Pcsk1* mRNA, at variable levels. All PMV neurons expressed very high *Pcsk1* mRNA levels, and the vast majority of DLL neurons expressed low to moderate *Pcsk1* mRNA levels. *Pcsk2* mRNA was expressed at high or very high levels by virtually all neurons in the same nuclei/regions.

### 2.6 Projections of PH/IFN Gcg neurons

To map the axonal projections of PH/IFN *Gcg* neurons, we injected Cre-dependent adeno-associated viruses (AAV) encoding a fluorescent reporter into the PH of *Gcg*-Cre mice. Besides intense Cre-dependent expression, low-level off-target (Cre-independent) reporter expression was detected by IF for all AAVs (**Fig S9** and *Methods*), a phenomenon reported before [46]. However, we observed no, or minimal off-target IF signal in axons. In addition, the modified IF protocol virtually eliminated the IF signal of off-target expressed enhanced green fluorescent protein (EGFP), while preserving the Cre-dependent signal (**Fig S9B-C1**). Therefore, we show EGFP IF with the modified protocol in AAV2/9-hSyn-DIO-hM3D(Gq)-EGFP-injected mice (**Fig 7**). The specificity of viral labeling was confirmed by injecting male *Gcg*-*Cre*^/*+*^;*tdTom*^/*+*^ mice with the same AAV. In the PH/IFN, the majority of EGFP-containing neurons expressed tdTomato (62.3 ± 2.6%, n=3), and dual-labeled EGFP/tdTomato fibers were distributed in the same pattern as described below (**Fig S10**).

**Figure 7.**
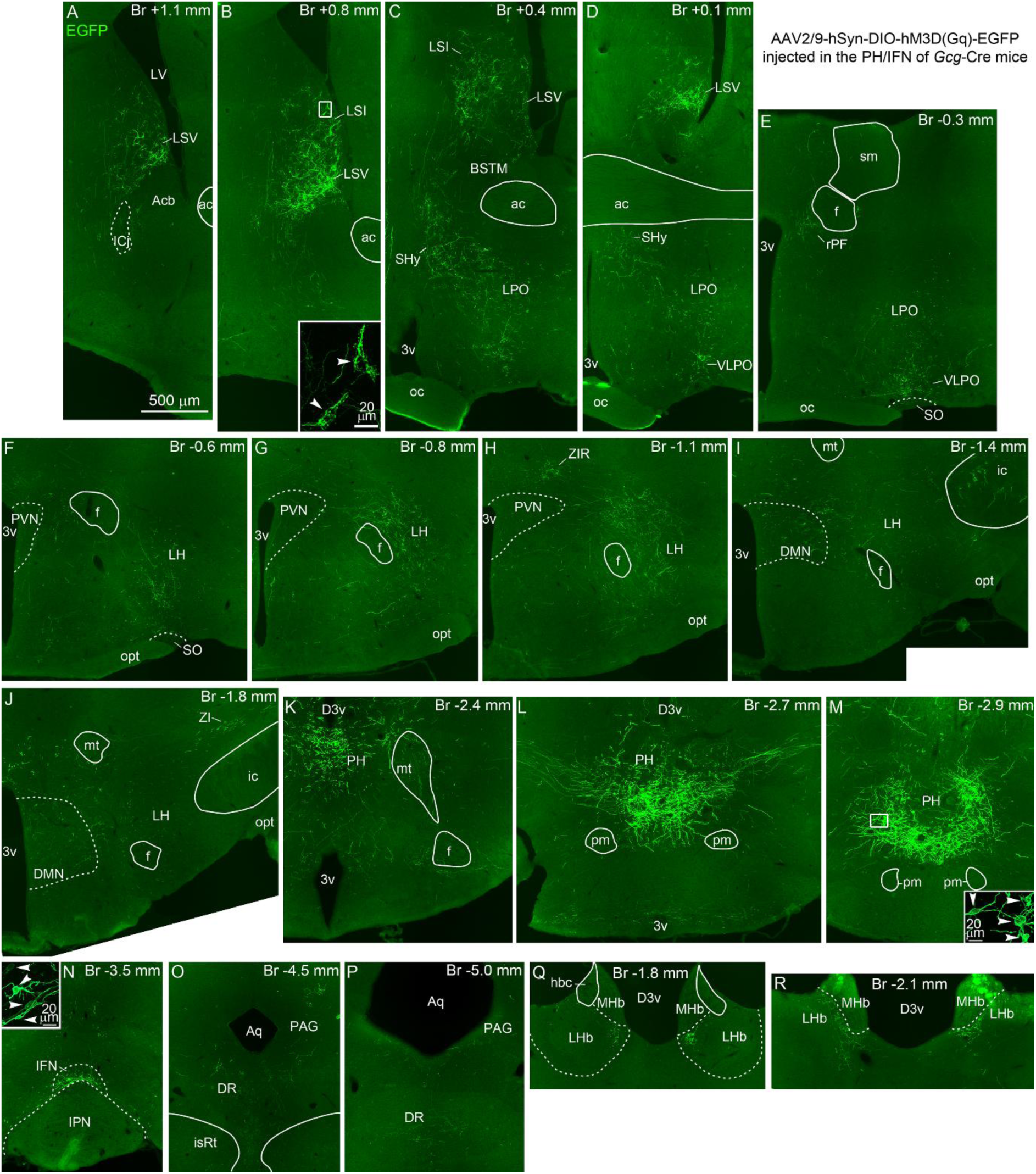
Projections of PH/IFN *Gcg* neurons shown by AAV-mediated tract-tracing. EGFP IF in a female *Gcg*-Cre^/Cre^ mouse injected with AAV2/9-hSyn-DIO-hM3D(Gq)-EGFP in the PH/IFN. Bregma levels are indicated in each panel. EGFP-labeled neurons in the PH are shown in **K**-**M**, and in the IFN in **N**. Insets in **M** and **N** show these cells (arrowheads) in high magnification confocal images. High or moderate densities of EGFP-labeled fibers are found in the ventral and intermediate lateral septum (**A-D**) where they form pericellular baskets (inset in **B**), preoptic region (**C-E**), ventrolateral preoptic nucleus (**D**, **E**), rostral perifornical area (**E**), lateral hypothalamus (**F-J**), zona incerta (**H**, **J**), hypothalamic dorsomedial nucleus (**I**, **J**), entopeduncular nucleus within the internal capsule (**I**), periaqueductal gray and dorsal raphe (**O**, **P**) and lateral habenula (**Q**, **R**). Inset in **B** shows pericellular baskets (arrowheads) formed by EGFP fibers in the lateral septum (high magnification confocal image of the boxed area). Abbreviations: 3v, third ventricle; ac, anterior commissure; Acb, nucleus accumbens; Aq, aqueduct; BSTM, medial part of the bed nucleus of the stria terminalis; D3v, dorsal third ventricle; DR, dorsal raphe; f, fornix; hbc, habenular commissure; ic, internal capsule; ICj, island of Calleja; IFN, interfascicular nucleus; IPN, interpeduncular nucleus; isRt, isthmic reticular formation; LH, lateral hypothalamus; LHb, lateral habenula; DMN, hypothalamic dorsomedial nucleus; LPO, lateral preoptic area; LSI, intermediate lateral septum; LSV, ventral lateral septum; LV, lateral ventricle; MHb, medial habenula; mt, mammillothalamic tract; oc, optic chiasm; opt, optic tract; PAG, periaqueductal gray; PH, posterior hypothalamic nucleus; pm, principal mammillary tract; rPF, rostral perifornical area; PVN, hypothalamic paraventricular nucleus; SHy, septohypothalamic nucleus; sm, stria medullaris; SO, supraoptic nucleus; VLPO, ventrolateral preoptic nucleus; ZI, zona incerta; ZIR, rostral zona incerta.

We analyzed 11 AAV-injected brains, both male and female (Supplementary Table 2). Injections that labeled neurons in the PH/IFN (7 brains) (**Fig 7K-N**), or only in the PH (2 brains) resulted in very similar fiber distributions. **Fig 7** shows overview images from a representative injection, **Fig S10** shows high magnification images of EGFP-labeled axons. The densest projections were observed in the ventral and intermediate lateral septum (**Fig 7A-D**), where fibers often formed pericellular baskets around cell bodies and dendrites (**Fig 7B** inset). Sparse fibers were observed in the dorsal lateral septum. In the nucleus accumbens, fiber density varied among mice from sparse to dense in two narrow zones, one adjacent to the Island of Calleja, the other lateral/ventrolateral to the lateral ventricle. The preoptic region, including the septohypothalamic nucleus and parts of the lateral and medial preoptic areas, received dense innervation (**Fig 7C**, **D**). Fibers were conspicuously dense in an area identified as the ventrolateral preoptic nucleus (VLPO) (**Fig 7D**, **E**), and moderately dense in the rostral perifornical area (**Fig 7E**). Dense projections were observed in the rostral half of the lateral hypothalamus, particularly dorsal to the supraoptic nucleus and dorsolateral/lateral to the fornix (**Fig 7F-H**), and in a confined area dorsal of the caudal part of the paraventricular nucleus, which we identified as part of the rostral zona incerta (**Fig 7H**). Moderate fiber density was observed in the caudal part of the lateral hypothalamus, the dorsomedial nucleus, the entopeduncular nucleus inside the internal capsule, and the lateral zona incerta (**Fig 7I**, **J**). Fibers were conspicuous in the medial part of the lateral habenula, particularly in its most caudal part (**Fig 7Q**, **R**). The only notable descending projection, with moderate fiber density, was observed in the PAG/DR (**Fig 7O**, **P**).

In two brains, AAV injections labeled both PH/IFN neurons and a significant number of PAG/DR neurons corresponding to the *Gcg* neuronal pattern (Supplementary Table 2). In these cases, dense projections were also observed in the oval nucleus of the bed nucleus of the stria terminalis and the central amygdala, known targets of PAG/DR neuron populations [47; 48].

#### 2.6.1 Axons of PH/IFN Gcg neurons contain PG/GLP-1

To examine whether PG/GLP-1 can be detected in fibers of AAV-labeled PH/IFN neurons, we performed dual or triple IF in a fed and two fasted females (Supplementary Table 2). In each mouse we detected light PG/GLP-1^C^ labeling in a small fraction of these fibers, mainly in pericellular baskets in the lateral septum, and occasionally in other areas, like the rostral zona incerta and lateral hypothalamus (**Fig 8A-C1**). These IF studies also visualized the projections of NTS/IRN *Gcg* neurons as intensely labeled PG/GLP-1^C^ fibers, alongside the EGFP- or enhanced yellow fluorescent protein (EYFP)-labeled axons of PH/IFN *Gcg* neurons. These two cell populations generally project to distinct but neighboring forebrain areas, with little overlap in the preoptic region, lateral hypothalamus, dorsomedial nucleus, lateral habenula and dorsal raphe (**Fig S11**). Even in the lateral hypothalamus, the two fiber systems were largely segregated to different subregions (**Fig S11F-H**).

**Figure 8.**
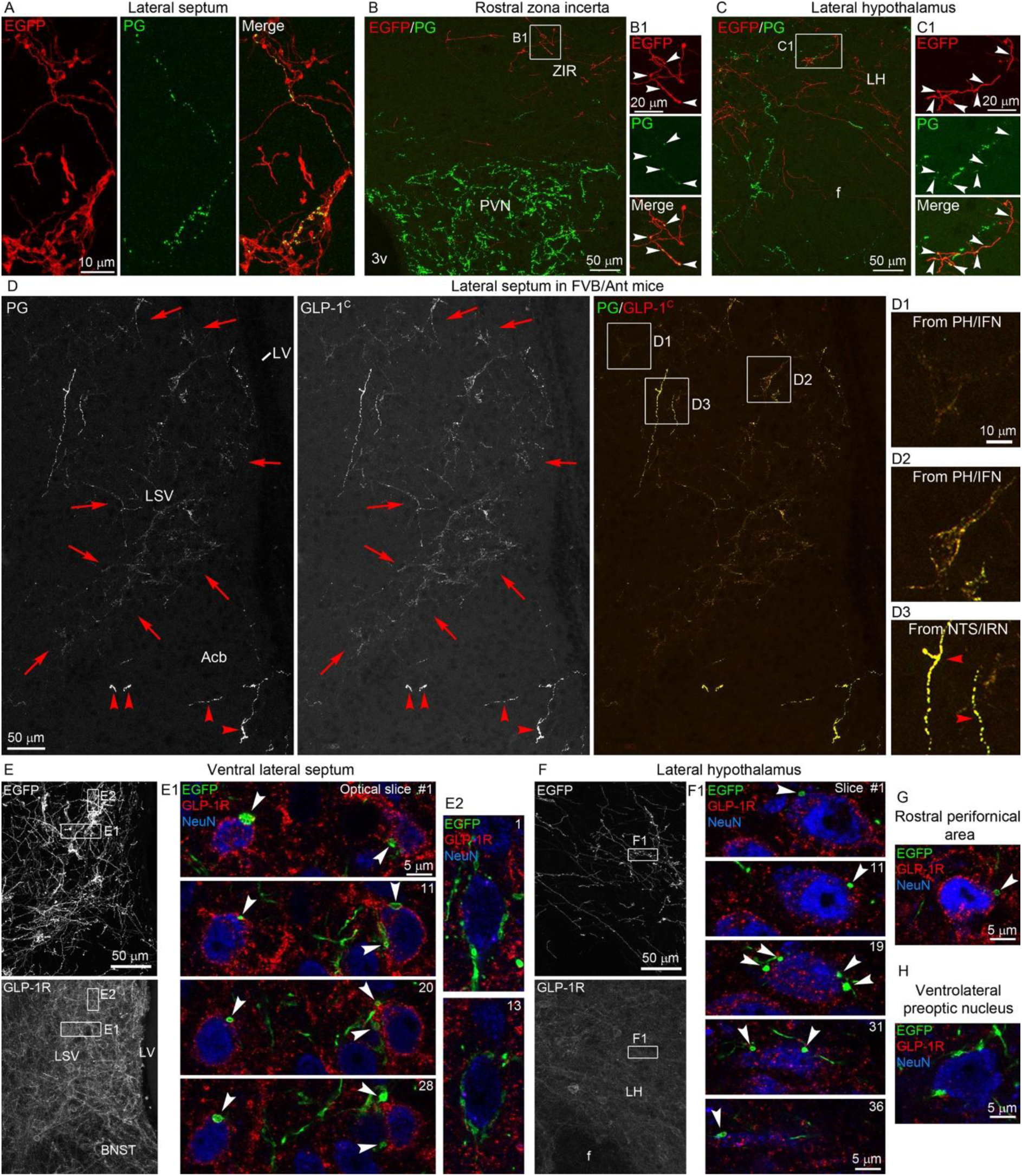
Axons of PH/IFN *Gcg* neurons contain PG/GLP-1 and innervate GLP-1R-expressing neurons. (**A**-**C**) Dual IF for EGFP (red) and PG (green) in a *Gcg*-Cre^/Cre^ mouse injected with AAV2/9-hSyn-DIO-hM3D(Gq)-EGFP in the PH/IFN shows PG-containing EGFP-labeled fibers in the lateral septum (**A**), rostral zona incerta (**B**), and lateral hypothalamus (**C**). The dual-labeled fibers inside the boxed areas in **B** and **C** are shown in higher magnification images (**B1**, **C1**, arrowheads). (**D**) Dual IF for PG/GLP-1^C^ in the lateral septum of an *ad libitum* fed male FVB/Ant mouse. Red arrows demarcate the area with lightly labeled, pericellular-basket-forming PG/GLP-1^C^ fibers. Red arrowheads in the nucleus accumbens point to intensely labeled PG/GLP-1^C^ fibers originating from the NTS/IRN. Magnified images of the boxed areas show pericellular baskets with light (**D1**) or medium-intensity labeling (**D2**) and intensely labeled axons from the NTS/IRN (**D3**, red arrowheads). (**E-H**) Triple IF for EGFP/GLP-1R/NeuN from a *Gcg*-Cre^/Cre^ mouse injected with AAV2/9-hSyn-DIO-hM3D(Gq)-EGFP in the PH/IFN. The monocolor images of the same fields show the distribution of EGFP-labeled fibers and GLP-1R in the lateral septum (**E**) and lateral hypothalamus (**F**). The boxed areas are shown in higher magnification color images (EGFP: green, GLP-1R: red, NeuN: blue). (**E1**) Four single optical sections (0.46 µm thickness, 0.23 µm Z-steps) show lateral septum GLP-1R neurons contacted by multiple EGFP-containing boutons (arrowheads). (**E2**) Two optical sections show a GLP-1R-negative neuron surrounded by boutons of a pericellular basket-forming EGFP-labeled axon. (**F1**) Five optical sections show several EGFP-labeled boutons (arrowheads) juxtaposed to the perikaryon and dendrite of a GLP-1R-positive neuron. (**G**) An EGFP bouton (arrowhead) is juxtaposed to a GLP-1R-positive neuron in the rostral perifornical area. (**H**) In the ventrolateral preoptic nucleus, EGFP-labeled boutons contact a GLP-1R-negative cell body. Abbreviations: Acb, nucleus accumbens; BNST, bed nucleus of the stria terminalis; f, fornix; LH, lateral hypothalamus; LSV, ventral lateral septum; LV, lateral ventricle; PVN, hypothalamic paraventricular nucleus; ZIR, rostral zona incerta.

In the lateral septum, the basket-forming axons of PH/IFN *Gcg* neurons could be easily distinguished from the sparse axons of medullary GLP-1 neurons, which are more intensely labeled for PG/GLP-1^C^ and do not form pericellular baskets. Therefore, we examined the lateral septum in male FVB/Ant mice for basket-forming PG/GLP-1^C^ axons (**Fig 8D-D3**). With dual IF, we observed such fibers in all mice, but their abundance was quite variable among brains. While we did not detect evidently more basket-forming PG/GLP-1^C^ fibers in fasted mice than in fed mice, their density largely correlated with the number of detected PG neurons in the PH/IFN. Of note, the PG and GLP-1^C^ signals completely co-localized in these axons (**Fig 8D-D2**). With IF, we detected neither GLP-1^N^ in PG-labeled basket-forming fibers, nor glucagon, using a glucagon-specific antibody (data not shown).

#### 2.6.2 PH/IFN Gcg neurons innervate both GLP-1R-positive and GLP-1R-negative neurons

To corroborate that PH/IFN *Gcg* neurons use GLP-1 as a neurotransmitter, we studied whether their axons target GLP-1R-expressing neurons. Using a specific antibody for GLP-1R, we performed triple IF for EGFP/GLP-1R/NeuN in two AAV2/9-hSyn-DIO-hM3D(Gq)-EGFP-injected *Gcg*-Cre mice, and looked for close appositions between EGFP-labeled fibers and GLP-1R-positive neurons. Additionally, we confirmed the presence or absence of *Glp1r* mRNA (encoding GLP-1R) expression in each projection site with FISH (**Fig S12**). We classified projection sites into three categories. The first category, where EGFP fibers frequently contacted GLP-1R-positive neurons, includes the lateral septum, lateral hypothalamus and rostral perifornical area. In the lateral septum and lateral hypothalamus, particularly dorsal/dorsolateral to the fornix, EGFP fibers often established few to multiple boutons on the perikarya or proximal dendrites of GLP-1R-positive neurons (**Fig 8E**, **E1**, **F**, **F1**). Fewer close appositions, 1-2 EGFP boutons per GLP-1R neuron, were observed in the rostral perifornical area (**Fig 8G**). In all three regions, EGFP fibers also made close contacts with GLP-1R-negative neurons. Most notably, the lateral septum neurons surrounded by pericellular basket-forming EGFP axons were GLP-1R-negative (**Fig 8E2**). In the second category, EGFP fibers overlapped with the distribution of GLP-1R neurons, but close appositions were rare. This category includes the preoptic region, dorsomedial nucleus, entopeduncular nucleus, lateral zona incerta and DR. The third category, where EGFP fibers did not overlap with GLP-1R neurons, includes the VLPO, rostral zona incerta, and lateral habenula. We confirmed the absence or paucity of *Glp1r* mRNA-expressing neurons in these areas (**Fig S12C**, **D**, **G**, **K**). In these projection sites, EGFP-labeled fibers frequently contacted the perikarya or main dendrites of GLP-1R-negative neurons, establishing few to multiple EGFP boutons on a single cell (**Fig 8H**).

## 3. Discussion

In this study, we systematically investigated the *Gcg*/PG system of the mouse brain and observed that *Gcg* neurons form 9 or 10 populations, distributed at various locations of the brain. A remarkable feature of this system is that one population, NTS/IRN *Gcg* neurons, expresses *Gcg* mRNA and PG protein at much higher levels than all other populations. Low *Gcg* expression levels are undoubtedly the main reason why so many *Gcg* neurons in other brain regions have remained undetected.

### 3.1 Comparison with previous studies

There are surprisingly few ISH studies on *Gcg* mRNA expression in the brain outside the medulla. In the first brain-wide study, Han and colleagues [4] identified medullary *Gcg* neurons with oligo DNA probes. Later, Merchenthaler and colleagues [8] detected low-level *Gcg* expression in the glomerular layer of the rat OB with radioactive riboprobes. Methodological differences likely explain why we identified more *Gcg* neuron populations than the latter study. For example, we used 10-fold lower RNase A concentration, which significantly increased detection sensitivity. Additionally, cells with very low mRNA levels are more noticeable in FISH-labeled sections than in film autoradiograms, which always have speckled background. Differences between rat and mouse *Gcg* expression patterns could also be a contributing factor. This may explain why we did not observe *Gcg* expression in the OB glomerular layer, although developmental or minimal expression in this layer was suggested in a BAC *Gcg*-iCre mouse model [18]. With RNAscope, Zheng and colleagues [16] observed *Gcg*-expressing cells in the granule cell layer of the OB, but not in the piriform cortex or BLA of adult rats, although they detected scattered *Gcg* neurons in the BLA of neonatal rats. However, the use of *Gcg*-Cre;tdTom animals, which have markedly reduced *Gcg* expression due to the IRES-iCre cassette insertion, might have contributed to the negative findings. A recent RNAscope study in mice by Ryu et al. [49] found scattered *Gcg-*expressing cells in several areas outside the medulla and OB. However, only occasional *Gcg* cells were detected in the PH/IFN, and none in the other populations we have identified. We suspect that the lower sensitivity of that study is partly due to the use of formalin-perfused brains that hinders mRNA detection [50].

*Gcg* promoter-driven transgene expression was often noted in locations corresponding to low-level *Gcg*-expressing neuron populations. Scattered cells were noted in the OB [27], piriform cortex [26; 27], PH [27], PMV [27], PAG/DR [6; 26; 27], and DLL [6]. A subpopulation of deep short-axon cells in the OB granule cell layer is transgenically labeled in preproglucagon-yellow fluorescent protein mice [17; 19; 20]. Large populations of Cre reporter positive neurons were observed in the OB, BLA and piriform cortex in both BAC *Gcg*-iCre mice [16; 18; 29], and knock-in *Gcg*-iCre rats [16]. In addition, a large number of reporter-positive cells appear in the piriform cortex, and probably the BLA, of knock-in *Gcg^iCre^* mice (see Figure 6d in Ref. [39]). Re-examining BAC transgenic *Gcg*-Cre mice [27], we observed that the Cre reporter tdTomato labeled a significant portion of *Gcg* neurons in the PH/IFN, PAG/DR, and OB granule cell layer. We observed minimal reporter expression in the piriform cortex and BLA, in contrast to other Cre-driver lines mentioned above. Thus, further studies on specific *Gcg* neuron populations will require the use of different Cre-driver strains.

### 3.2 The brain PG system: peptidergic nature and functional implications

Our IF studies demonstrate that *Gcg* mRNA is translated into PG protein in all *Gcg*-expressing neuron populations. Although in most regions we detected much fewer PG-containing neurons than *Gcg* mRNA-expressing neurons, this can be explained with the lower sensitivity of IF compared to FISH. Further immunohistochemical studies employing signal amplification techniques might detect significantly more PG neurons. We also observed GLP-1^C^ immunolabeling in a subset of PG neurons in all populations except the piriform cortex. This observation, and that all *Gcg* neuron populations express *Pcsk1*, suggest that most, if not all, *Gcg* neurons can produce GLP-1. The very sporadic labeling of the GLP-1^N^ antibody is likely due to its lower affinity, or lower levels of bioactive GLP-1 than the C-terminally amidated GLP-1 forms, and not because of a lack of bioactive GLP-1. GLP-1 production in multiple brain regions largely explains why GLP-1R is so widely distributed in the brain [8; 51–54]. After the initial observation that *Glp1r* is expressed more broadly than the projection sites of medullary *Gcg* neurons [8], discussions on this topic have focused mainly on the ventral hippocampus and cortex [14; 55; 56]. In fact, *Glp1r* is expressed in numerous areas with little to no innervation from medullary *Gcg* neurons, including the striatum, amygdalohippocampal area, mammillary nuclei, inferior and superior colliculus, and others (see overview images of *Glp1r*-GFP distribution in Supplementary Figure 6 of Ref. [57], which closely correspond to our *Glp1r* FISH data). While it remains to future studies to map the projections of each *Gcg* neuron population, known connectivity patterns suggest GLP-1-containing inputs to some of the areas mentioned. For example, GLP-1 innervation of the ventral hippocampus might originate from the BLA [58]. Cortical GLP-1R neurons likely receive GLP-1-containing innervation from the claustrum, which projects almost exclusively to the cerebral cortex [59]. GLP-1R neurons in the inferior colliculus probably receive GLP-1 input from the DLL [60].

GLP-1 is produced simultaneously with GLP-2 [40], and the two peptides are likely co-released [61; 62]. However, GLP-2 receptor (GLP-2R) distribution in the brain, while also widespread, differs significantly from that of GLP-1R [24; 61; 63–66]. It remains an intriguing question whether a single *Gcg* neuron population may regulate distinct downstream neuron populations selectively *via* GLP-1R or GLP-2R signaling. We also show that all *Gcg* neuron populations express *Pcsk2*, suggesting that they can produce glucagon. With IF we observed glucagon signal in the majority of the axons of medullary GLP-1 neurons, confirming their glucagon synthesis (data not shown). Whether glucagon functions as a neuropeptide depends also on the neuroanatomical distribution of the glucagon receptor, which remains largely unknown, but appears more restricted than GLP-1R or GLP-2R distribution [64; 67; 68]. Species differences may also exist, as glucagon levels were reported to be higher than GLP-1 levels in the canine brain [69].

*Gcg* is expressed in a peculiar collection of brain nuclei with diverse functions. The OB and piriform cortex are part of the olfactory system [70], the claustrum modulates cortical activity [59], the BLA is involved in fear learning and reward reinforcement [71], the hippocampus is essential in memory formation and spatial navigation [72], the PAG/DR regulates various types of behavior [73], the DLL is part of the auditory brainstem [74], and the PMV in male mice regulates social aggression [75; 76]. We hypothesize that *Gcg* might be an activity-dependent gene in some of these populations, given its varying expression levels particularly among cells of the posterior hippocampus and piriform cortex, but also its fasting-induced upregulation in the PH/IFN. *Gcg* might be also expressed in additional brain regions under certain conditions.

### 3.3 PH/IFN Gcg neurons

After medullary *Gcg* neurons, we observed the second highest *Gcg* and PG expression levels in the PH/IFN of fasted mice. Based on our cell counts from wild-type and *Gcg*-Cre;tdTomato mice, we estimate the size of this population between 50-75% of the number of NTS/IRN *Gcg* neurons. The finding that fasting upregulates *Gcg* mRNA, PG and GLP-1 synthesis in the PH/IFN is surprising and unexpected. First, despite the anorexigenic nature of GLP-1, PH/IFN *Gcg* neurons are regulated similarly to orexigenic agouti-related protein (AGRP) neurons in the arcuate nucleus, in which fasting markedly increases AGRP and neuropeptide Y expression [77; 78]. We also have preliminary data that fasting induces c-fos expression, a marker of cellular activation, in PH/IFN *Gcg* neurons [79], similar to AGRP neurons [80; 81]. Second, feeding status regulates PH/IFN *Gcg* neurons oppositely to medullary *Gcg* neurons: *Gcg* expression in the medulla remains unchanged or decreases during fasting [82; 83], and medullary *Gcg* neurons become active in response to feeding [84–87]. Third, the PH and IFN are known to mediate physiological and behavioral stress responses [88–91], but not parts of the known feeding-related circuits.

How PH/IFN *Gcg* neurons may contribute to the adaptation to food deprivation remains to be investigated. PH/IFN *Gcg* neurons project to several areas involved in the motivational or reward aspects of feeding behavior, including the ventral lateral septum, lateral hypothalamus and lateral habenula [92–95]. For example, distinct lateral hypothalamic neuron groups modulate taste preferences *via* projections to the lateral septum or the lateral habenula [94]. PH/IFN *Gcg* neurons may modulate this or similar neuronal circuits that regulate the motivation to eat, including foraging. Interestingly, only a subset of the downstream neurons of PH/IFN *Gcg* neurons are GLP-1-receptive, located mainly in the lateral septum and lateral hypothalamus. Other target neurons lacking GLP-1R may be regulated *via* a fast synaptic neurotransmitter, other neuropeptides, or indirectly by GLP-1 facilitating the activity of their GLP-1R-expressing synaptic inputs [96].

### 3.4 Conclusions

The *Gcg* system of the mammalian brain is much more complex than previously understood. Uncovering the functions of this system in animal models, and its anatomy in the human brain are increasingly timely, as clinically approved GLP-1 analogues may penetrate the brain in sufficient amounts to mimic the functions of GLP-1-containing neural circuits.

## 4. Methods

### 4.1 Animals

Mice were group housed (2-5 per cage) on corn cob bedding under standard environmental conditions (lights on between 06:00 and 18:00 h, temperature 23 ± 1 °C, mouse chow and water *ad libitum*). All experiments followed the ARRIVE guidelines and were carried out in accordance with EU Directive 2010/63/EU on the protection of animals used for scientific purposes. Experimental protocols were reviewed and approved by the Animal Welfare Committee at the HUN-REN Institute of Experimental Medicine (HUN-REN IEM) and the Hungarian Scientific Ethics Committee for Animal Experimentation (PE/EA/1102-7/2020, PE/EA/00519-6/2025).

Wild-type mice from the C57BL/6J and FVB/Ant strains were used from the local colony of HUN-REN IEM. Transgenic *Gcg*-Cre mice were generated on the C57BL/6J background and express Cre recombinase from the *Gcg* locus of a BAC [27]. The line was rederived at HUN-REN IEM, as described previously [97]. *Gcg*-Cre;tdTomato mice were generated by crossing *Gcg*-Cre mice with Ai9 mice (strain # 007909, The Jackson Laboratory; C57BL/6J genetic background) carrying a knock-in transgene at the Gt(ROSA)26Sor locus in which a loxP-flanked STOP cassette prevents transcription of CAG promoter-driven tdTomato. In Cre-expressing cells, Cre-mediated removal of the STOP cassette results in tdTomato expression.

### 4.2 Fasting experiments

At the beginning of the light phase (06:00-07:00 h), group-housed littermates (4-5 per cage) were divided into two groups and placed in separate cages (2-3 per cage) with clean corn cob bedding. Chow was withheld from the fasted group and provided *ad libitum* to the control group. All mice were euthanized 30h later. In each fed *vs* fasted comparison, both control and fasted groups (n=4-5 mice) included mice from 2-3 different litters. In the 48h fasting experiment, mice were housed individually when the experiment began.

### 4.3 Tissue collection for FISH

Mice were anesthetized with isoflurane and decapitated. The brains were removed and snap-frozen in powdered dry ice. Coronal 16 μm thick sections were cut on a Leica CM3050 S cryostat (Leica Microsystems), thaw-mounted on Superfrost Plus glass slides (Epredia), and air-dried. Sections were collected in one-in-twelve series (192 µm distance between two consecutive serial sections) from the whole brain or select regions (PH/IFN, etc.) and stored at -80°C until FISH.

### 4.4 FISH riboprobes

The following digoxigenin-labeled antisense riboprobes were used (NCBI GenBank accession numbers in parentheses): *Gcg*, bases 75-944 of murine *Gcg* mRNA (NM_008100.4) [30]; *Glp1r*, 11-1402 of murine *Glp1r* mRNA (NM_021332.2) [30]; *Pcsk1*, 287-1286 of murine *Pcsk1* mRNA (NM_013628.3); *Pcsk2*, 431-1430 of murine *Pcsk2* mRNA (NM_008792.4). To detect *tdTomato* mRNA, we used a fluorescein-labeled antisense riboprobe corresponding to the red fluorescent protein (RFP) sequence, which is in tandem dimer form in the synthetic *tdTomato* gene (22-677 and 748-1403 of AY678269.1). *Pcsk1, Pcsk2,* and *tdTomato* template DNAs were synthesized and cloned into pBluescript II vectors by GenScript Biotech.

### 4.5 FISH combined with IF

Sections were hybridized with one of the digoxigenin-labeled riboprobes. The FISH procedure followed the protocol previously described for fresh frozen sections [50], except that 2 µg/ml RNase A was used in the post-hybridization phase, instead of 20 µg/ml, which was critical to increase detection sensitivity. To detect the digoxigenin-labeled probes, sections were incubated overnight in peroxidase-conjugated sheep anti-digoxigenin antibody Fab fragments (Roche, Cat# 11207733910), diluted at 1:100 in 1% blocking reagent for nucleic acid hybridization (Roche, Cat# 11096176001). Signal amplification was performed with the TSA Plus Biotin Kit (Akoya Biosciences, Cat# NEL749A001KT) for 30 min, using the biotin reagent at 1:500 dilution in 0.05M Tris (pH 7.6) containing 0.01% H_2_O_2_. Biotin deposits were detected with Alexa Fluor 488-conjugated Streptavidin (ThermoFisher, 1:500). Sections were incubated overnight in a Guinea pig antiserum against NeuN (1:2,000), and then in Cy3-conjugated anti-Guinea pig IgG (Jackson Immunoresearch, 1:200) for 2h. Sections were coverslipped with SlowFade Diamond mountant containing DAPI (ThermoFisher).

### 4.6 Dual-label FISH

Sections were hybridized with a cocktail containing the fluorescein-labeled *tdTomato* riboprobe and a digoxigenin-labeled riboprobe for either *Gcg*, *Pcsk1* or *Pcsk2*. The procedure followed the above protocol, except the final detection steps. Sections were first incubated in peroxidase-conjugated sheep anti-digoxigenin Fab fragments overnight, and the hybridization signal was amplified with the TSA Plus DIG Kit (Akoya Biosciences, Cat# NEL748E001KT) for 30 min, applying the digoxigenin amplification reagent at 1:500 dilution in 0.05M Tris containing 0.01% H2O2. Sections were incubated in a rabbit monoclonal anti-digoxigenin antibody (ThermoFisher, Cat# 700772; 1 μg/ml) for 2 h, in the presence of 2% sodium azide to inactivate peroxidase activity. Sections were rinsed in PBS and incubated overnight in peroxidase-conjugated sheep anti-fluorescein Fab fragments (Roche, Cat# 11426346910), diluted 1:100 in 1% blocking reagent. The *tdTomato* hybridization signal was amplified with the TSA Plus Biotin Kit for 30 min. Sections were incubated in the cocktail of Alexa Fluor 488-conjugated Streptavidin and Alexa Fluor 555-conjugated donkey anti-rabbit IgG (ThermoFisher; 1:500) for 2h. Native tdTomato fluorescence was completely abolished by the FISH conditions.

### 4.7 AAV injection into the PH/IFN

*Gcg*-Cre or *Gcg*-Cre;*tdTomato* mice (7-15 wks) were anesthetized ip with ketamine-xylazine (ketamine: 50 mg/kg; xylazine: 10 mg/kg body weight) and their head positioned in a stereotaxic apparatus. Through a burr hole in the skull, a glass pipette (25 μm inner tip diameter) connected to a Nanoject III injector (Drummond Scientific Company) was lowered into the brain at stereotaxic coordinates corresponding to the PH (anteroposterior: -2.54 mm, mediolateral: -0.2 mm, dorsoventral: -4.75 mm [35]). Mice were injected with 400 nl (10 nl/sec) of one of the following AAVs: AAV1-EF1a-DIO-hM3D(Gq)-mCherry (titer: 7×10^12^/ml, or its 10-fold dilution; Duke University Viral Vector Core), AAV8-EF1a-DIO-hM3D(Gq)-mCherry (9×10^12^/ml; Duke), AAV2/9-hSyn-DIO-hM4D(Gi)-EYFP (>2×10^12^/ml, BrainVTA, Wuhan, China), AAV2/9-hSyn-DIO-hM3D(Gq)-EGFP (>2×10^12^/ml, BrainVTA), and AAV8-hSyn-DIO-hM3D(Gq)-mCherry or AAV8-hSyn-DIO-hM4D(Gi)-mCherry (5.3-5.7×10^12^/ml, University of North Carolina Vector Core). Three minutes after the injection, the pipette was slowly removed and the wound closed. Mice were euthanized 4-6 weeks later by transcardial perfusion. To increase Cre expression, and thus the number of AAV-labeled neurons, some mice were fasted for 24h, 1-2 weeks after AAV injection. To facilitate detection of PG/GLP-1 in AAV-labeled fibers, two mice were fasted for 30h directly before perfusion.

### 4.8 Tissue collection for IF and direct tdTomato fluorescence

Mice were anesthetized with ketamine-xylazine and perfused transcardially with 10 ml PBS (pH 7.4), followed by 40 ml 4% paraformaldehyde in 0.1 M phosphate buffer (PB; pH 7.4). The brains were removed, postfixed in 4% paraformaldehyde for 2 h, then cryoprotected in 30% sucrose in PBS overnight. Brains were snap-frozen on dry ice and cut on a cryostat or freezing microtome (Leica Microsystems) into one-in-four series of 25 µm thick coronal sections. Sections were stored in anti-freeze solution (30% ethylene glycol, 25% glycerol, 0.05 M PB) at -20°C until used. To observe direct tdTomato fluorescence, sections from Gcg-Cre;tdTomato brains were briefly washed in 0.05 M Tris before mounted on glass slides and examined with an epifluorescent microscope.

### 4.9 Standard IF

Our standard IF protocol included a permeabilization step of 0.5% Triton X-100 and 0.5% H2O2 in PBS for 20 min, rinses in PBS (3×10 min), and a 20 min blocking step with antibody diluent (2% normal horse serum, 0.2% Kodak Photo-Flo, 0.2% sodium azide in PBS) to reduce non-specific antibody binding. The sections were incubated in primary antibodies for 16-20h at room temperature, rinsed with PBS, then incubated in fluorochrome-conjugated donkey secondary antibodies, for 3-4 h. This protocol was applied to detect mCherry in brains injected with an mCherry-encoding AAV, with a sheep tdTomato antiserum (1:80,000), and Alexa Fluor 555-conjugated anti-sheep IgG (ThermoFisher, 1:500). To triple-label for EGFP/GLP-1R/NeuN in AAV-injected brains, we used a sheep YPet antiserum (1:20,000) to detect EGFP, a rabbitized monoclonal antibody against GLP-1R (0.025 µg/ml) and the Guinea pig NeuN antiserum (1:4000). The applied secondary antibodies were Alexa Fluor 488-conjugated anti-sheep, Alexa Fluor 555-conjugated anti-rabbit (ThermoFisher, 1:500 each) and Cy5-conjugated anti-Guinea pig (Jackson, 1:300) IgGs. We initially used this protocol for IF with the PG antibody (0.2 µg/ml) and Alexa Fluor 555-conjugated anti-rabbit IgG.

### 4.10 Modified IF

Use of the mouse GLP-1 antibodies with fluorochrome-conjugated anti-mouse IgGs by the standard IF protocol resulted in unacceptably high background in all brain regions, but particularly in circumventricular organs due to the entry of mouse IgGs [98]. Stemming from our observation that ISH conditions disrupt the antigenicity of endogenous IgGs, we first incubated the sections in ISH buffer (composition described in Ref. [50]) at 58 ^0^C for ∼20h and then proceeded with the standard protocol. This step eliminated the background hindering GLP-1 detection (**Fig S4C-G**), enhanced the PG and GLP-1 signals (**Fig S4A-G**) due to antigen unmasking [99], and completely or substantially reduced lipofuscin and other autofluorescence.

For PG/GLP-1^C^/NeuN IF, the rabbitized PG antibody (0.2 µg/ml), the mouse GLP-1^C^ antibody (1.0 µg/ml), and the Guinea pig NeuN antiserum (1:4,000) were used, with Alexa Fluor 488-conjugated anti-rabbit, Alexa Fluor 555-conjugated anti-mouse (ThermoFisher, 1:500) and Cy5-conjugated anti-Guinea pig IgGs (Jackson, 1:300). For PG/GLP-1^N^/NeuN IF, sections were first incubated in the mouse GLP-1^N^ antibody (0.2 µg/ml), then in the cocktail of PG and NeuN antibodies, and finally in the cocktail of the same secondary antibodies. For PG/NeuN IF in the BLA of female C57BL/6J mice, the primary antibodies were detected with Alexa Fluor 555-conjugated anti-rabbit and Cy5-conjugated anti-Guinea pig IgGs. For PG/glucagon IF, we used a mouse anti-glucagon antibody (0.05 µg/ml) in cocktail with the PG antibody, and Alexa Fluor 488-conjugated anti-rabbit and Alexa Fluor 555-conjugated anti-mouse IgGs. For PG/tdTomato/NeuN labeling in *Gcg*-Cre;tdTomato mice, we applied the sheep tdTomato antiserum (1:80,000) in cocktail with the PG and NeuN antibodies, with Alexa Fluor 488-conjugated anti-rabbit, Alexa Fluor 555-conjugated anti-sheep, and Cy5-conjugated anti-Guinea pig IgGs. For EGFP/PG/GLP-1^C^ or EYFP/PG/GLP-1^C^ labeling in AAV-injected *Gcg*-Cre mice, we used the sheep YPet antiserum (1:20,000) to detect EGFP or EYFP, in cocktail with the other primary antibodies, and Alexa Fluor 488-conjugated anti-sheep, Alexa Fluor 555-conjugated anti-rabbit, and Alexa Fluor 647-conjugated anti-mouse IgG (Jackson Immunoresearch, 1:200). For EGFP/tdTomato IF in AAV-injected *Gcg*-Cre;tdTomato mice, we used a rabbit anti-RFP antibody (1:1500) for tdTomato in cocktail with the sheep YPet antiserum, and Alexa Fluor 488-conjugated anti-sheep and Alexa Fluor 555-conjugated anti-rabbit IgGs. Sections were coverslipped with SlowFade Diamond.

### 4.11 Antibody specificity

The Guinea pig NeuN antiserum (Cat# ABN90, Millipore) labeled the characteristic neuronal pattern in the brain, labeling both the nucleus and cytoplasm [30].

To detect PG, we selected the best-performing antibody from our preliminary tests (Supplementary Table 3). The PG antibody is a chimeric rabbitized version of a mouse monoclonal antibody (clone 62-2F6, Novo Nordisk A/S [30; 100]) that binds to the GLP-1 (12-22) epitope, corresponding to PG 83-93. Such midportion GLP-1 antibodies tend to recognize the epitope in all precursors [43; 101]. We confirmed this by preincubating the working antibody dilution with an N- and C-terminally extended peptide, PG 76-153 (ThermoFisher, Cat# RP-102746; 20 µg/ml of His-ABP-tagged peptide), which resulted in the absence of immunolabeling (**Fig S3A**, **B**). Despite epitope homology with the glucagon (6-16) sequence, the antibody does not cross-react with glucagon, as preincubation with 5 µg/ml glucagon (1-29) (Genscript, Cat# RP10772) did not affect the immunolabeling. This antibody appears completely specific to PG in the mouse brain, based on the various co-localization studies (see *Results*) and that PG-labeled cell bodies were detected exclusively in areas where *Gcg* mRNA is expressed. The chimeric version of the antibody was generated by grafting the antigen-binding domains of the mouse antibody onto rabbit IgG constant domains [54].

The mouse monoclonal GLP-1^C^ antibody (HYB 147-06, clone ID: 8G9; Statens Serum Institut, Denmark) recognizes all forms of GLP-1 with amidated C-terminus. Preincubating the antibody with the PG 76-153 peptide did not affect the immunolabeling (**Fig S3C**, **D**). The GLP-1^C^ signal always co-localized with the PG antibody signal.

The mouse monoclonal GLP-1^N^ antibody (NBP2-23558, Novus Biologicals) binds the free N-terminus of GLP-1 (7-36)amide and GLP-1 (7-37), and shows <0.2% cross-reactivity with GLP-1 (1-37), and ∼1% with GLP-2. Pre-incubating the antibody with the PG 76-153 peptide did not significantly affect the immunolabeling (**Fig S3E**, **F**). The free N-terminus epitope partially overlaps with the PG antibody epitope, as applying the GLP-1^N^ antibody in cocktail with, or following the PG antibody resulted in only PG but no GLP-1^N^ labeling. Dual PG/GLP-1^N^ IF was achieved by incubating the sections first in GLP-1^N^, then in PG antibody.

The mouse monoclonal glucagon antibody (clone K79bB10, Cat# SAB4200685; Sigma-Aldrich) binds to glucagon (1-29) and exhibits weak cross-reactivity with oxyntomodulin. Its specificity for IF was demonstrated by the absence of immunolabeling in alpha-cell specific *Pcsk2* knockout pancreatic islets, which lack glucagon but contain other PG-derived peptides, including oxyntomodulin [102].

The sheep tdTomato antiserum generated in our laboratory [103], and the rabbit RFP antibody (Cat# 600-401-379, Rockland) recognize tdTomato and mCherry and show no immunolabeling in wild-type mouse brains.

A sheep antibody against the YFP variant YPet was generated in-house. The YPet coding region was inserted into the pET26b (+) bacterial expression vector (Merck) by adding a C-terminal His-tag. Recombinant expression in Rosetta 2(DE3) *E. Coli* strain (Merck), and isolation of His-tagged YPet was performed as described previously [103]. For immunization, 760 μg YPet in 1 ml PBS was emulsified with an equal volume of Freund’s complete adjuvant (Sigma-Aldrich) and injected intracutaneously into sheep. Subsequent boosts with Freund’s incomplete adjuvant were administered at 28-day intervals. Eight days after the fourth immunization, blood was collected and the serum was separated by centrifugation. The serum was affinity-purified by YPet-containing cyanogen bromide-activated Sepharose gel. This antiserum recognizes YPet, EGFP and EYFP, and yields no immunolabeling in the brains of wild-type mice.

The specificity of the GLP-1R antibody (clone 7F38, Novo Nordisk A/S) was previously demonstrated by the lack of immunolabeling in GLP-1R deficient mice in immunohistochemical studies [52; 54]. GLP-1R IF signal was observed along the cell membrane and intracellularly, concentrating in the Golgi compartment **(Fig 8E1**, **F1**, **G**), as previously described [30].We used the rabbitized version of the mouse monoclonal antibody [54].

### 4.12 Imaging

Low magnification fluorescent images were captured with the 10× objective of a Zeiss Axio Imager M2 epifluorescent microscope equipped with AxioCam MRc5 and AxioVision Se64 Rel.4.9.1 software. Higher magnification confocal images (Z-stacks) were taken with a Zeiss LSM 780 or LSM 900 confocal microscope, using Plan-Apochromat 20× and 63× lenses. Line-by-line sequential scanning was performed with standard laser lines (488, 561, 633 or 640 nm) and beam splitters (MBS 488/561/633 or 488/561/640). In LSM 780, detection wavelengths were set for 499-552 nm for Alexa 488, 570-623 nm for Alexa 555/Cy3, and 638-755 nm for Alexa 647/Cy5. In LSM 900, detection wavelengths were 410-545 nm for Alexa 488, 545-620 nm for Alexa 555/Cy3, and 656-700 nm for Alexa 647/Cy5. Z-projections of optical slices were made with Zen software (Zeiss). Adobe Photoshop (Adobe Inc) was used to create composite images and to modify brightness and contrast for illustrations.

### 4.13 Cell counts, image analysis and statistics

Cell counts were conducted manually with the epifluorescent microscope. *Gcg* neurons were counted bilaterally, in every 12th 16 µm thick section through the brain for whole-brain mapping, or through the PH/IFN in fasting experiments. Background levels varied from essentially none to scattered fluorescent dots. Higher background levels occasionally occurred toward section edges, rarely affecting the piriform cortex or claustrum. Generally, we counted a neuron *Gcg*-positive if at least 3 hybridization dots were present in a NeuN-positive cell profile. This criterion was impractical for the piriform cortex and OB, where determining the associations of a few (3-5) hybridization dots between densely packed cell bodies was difficult without confocal imaging. For these two areas, we considered clear hybridization dot clusters as *Gcg* positive neurons. In the dual-label FISH studies, cells were also counted in every 12th 16 µm thick section through a specific *Gcg-*expressing neuron population. Here we regarded a single hybridization dot in a *tdTom* cell as positive for *Gcg*, *Pcsk1* or *Pcsk2* mRNA. In IF, direct tdTomato fluorescence and AAV-injection studies, cells were counted bilaterally in every 4th 25 µm thick section through the brain or a specific *Gcg*-expressing nucleus.

*Gcg* FISH signal was quantified with ImageJ software (https://imagej.net/ij/) in images captured with the 10× objective of the epifluorescent microscope. The signal was separated from the background using the same threshold value for all images, and integrated density was measured.

Close contacts between EGFP fibers and GLP-1R-positive neurons were examined on confocal Z-stacks captured with the 63× objective (0.46 µm thick optical slices, 0.23 µm Z-steps). We examined at least 2 different fields of view for each projection area per animal.

Paxinos and Franklin’s atlas [35] was used to indicate anteroposterior (Bregma) levels and for anatomical nomenclature, except for cortical areas and the hippocampus, where we followed the Allen Brain Atlas [34].

GraphPad Prism 10 software was used to calculate statistics and plot graphs. All data are presented as mean ± SEM. Two-way or one-way ANOVA, followed by Tukey’s multiple comparisons test, were used to compare the effects of fasting on *Gcg* expression across sexes and strains. Two-tailed unpaired t-test was used to compare PG neuron numbers between fed and fasted mice. Differences were considered significant if p < 0.05.

## Author contributions

GW: Conceptualization, Formal Analysis, Investigation, Methodology, Validation, Visualization, Writing - original draft, Writing - review and editing. AK: Investigation, Validation, Writing - review and editing; PM: Resources; MGR: Resources; YR: Investigation; IV: Investigation; BD: Investigation, Resources; AH: Resources; ZL: Resources; BG: Resources. CF: Funding Acquisition, Resources, Supervision, Writing - review and editing.

## Acknowledgements

This study was funded by the National Brain Research Program (NAP 3.0; NAP2022-I-10/2022) of the Hungarian Academy of Sciences and the National Research, Development and Innovation Office (NKFIH Advanced grant N. 150936 and N. 153242), and supported by the National Academy of Scientist Education Program of the National Biomedical Foundation. The authors thank Dr. Joel Elmquist (UT Southwestern Medical Center, Dallas, TX, USA) and Dr. Michael M. Scott (University of Virginia, Charlottesville, VA, USA) for the kind donation of the *Gcg*-Cre mouse line. The authors thank Ágnes Simon for her expert technical assistance.

