## Supplementary Material for "Brain-wide mapping of proglucagon expression in mice identifies fasting-responsive GLP-1 neurons in the posterior hypothalamic nucleus"

### Supplementary Figures and Tables

### Figure S1

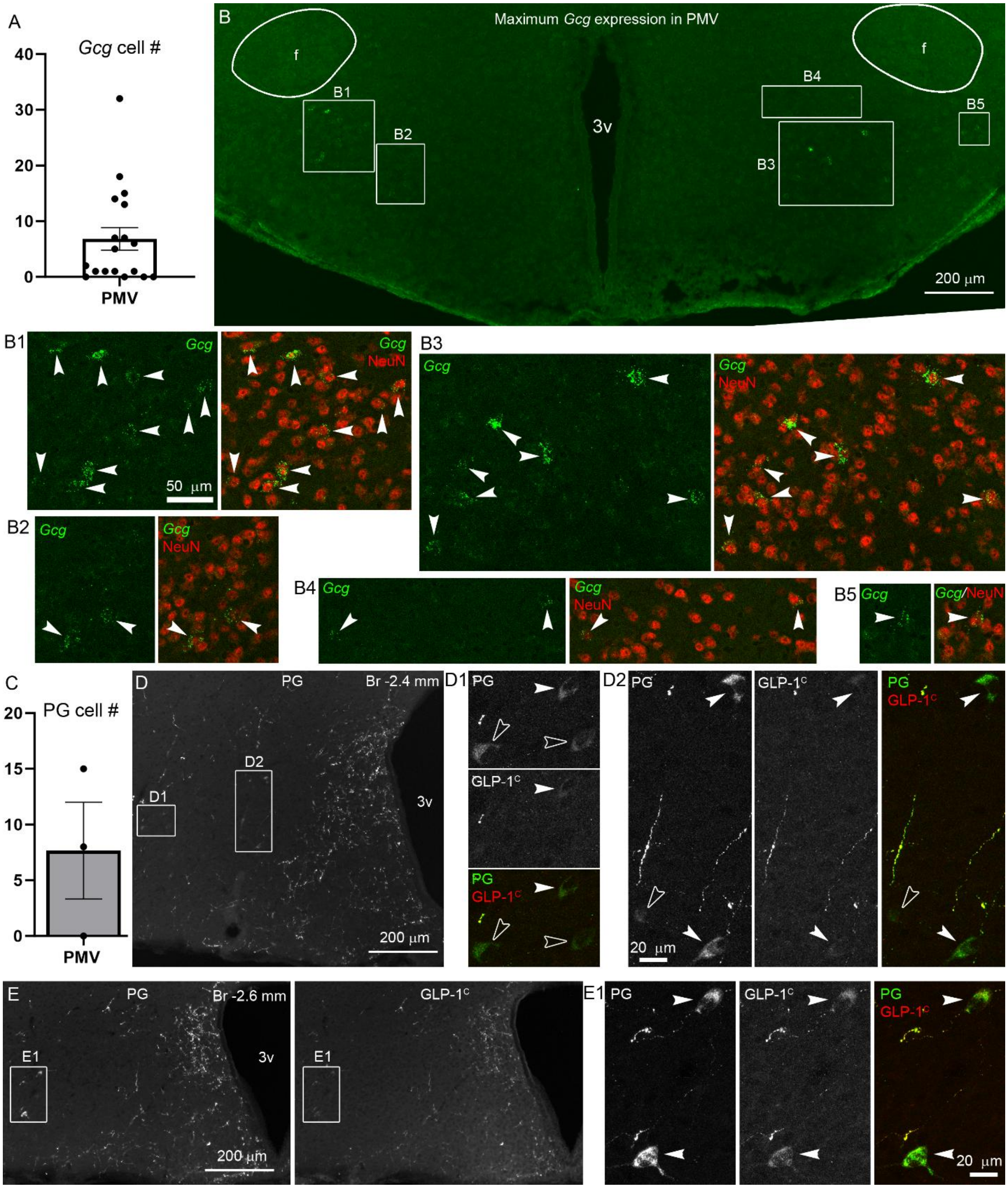


**Figure S1. *Gcg* mRNA, PG and GLP-1 expression in the ventral premammillary nucleus (PMV) of male C57BL/6J mice.** (**A**) Graph showing the number of *Gcg* neurons counted in the PMV and its vicinity (counted in a series of every 12th 16 µm thick section, in which 2 or 3 sections contained the PMV) in 18 male C57BL/6J mice (11-20 wks; 10 *ad libitum* fed, 8 fasted). (**B**) Maximum *Gcg* expression detected in the PMV by FISH. Higher magnification confocal images of the boxed areas (**B1**-**B5**) show *Gcg* mRNA-expressing neurons (arrowheads). The merged *Gcg* (green)/NeuN (red) images show that the *Gcg* FISH signal is localized in neuronal perikarya. (**C**) Graph showing the number of PG neurons in the PMV, detected by IF, in 3 male C57BL/6J mice (counted in a series of every 5th 20 µm thick section, in which 3-4 sections contained the PMV). (**D**, **E**) Dual-label IF for PG/GLP-$1^C$ in the PMV of a male C57BL/6J mouse. (**D**) Low magnification image shows the PG signal in the PMV. Higher magnification confocal images of the boxed areas (**D1**, **D2**) show PG, GLP-$1^C$, and merged PG/GLP-$1^C$ signals. Arrowheads point to PG neurons lightly positive for GLP-$1^C$. Open arrowheads indicate GLP-$1^C$-negative PG neurons. (**E**) Low magnification images shows the PG and GLP-$1^C$ signals in the PMV in separate panels. Higher magnification confocal images of the boxed area (**E1**) show two PG neurons with moderately intense GLP-$1^C$ signal (arrowheads). Abbreviations: 3v, third ventricle; f, fornix.

**Figure S2**

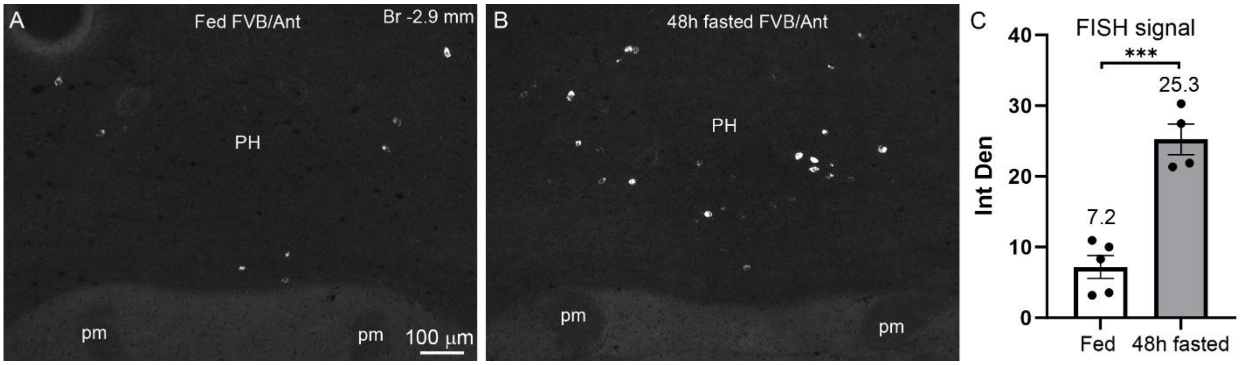


**Figure S2. Effect of 48h fasting on *Gcg* expression in the PH of male FVB/Ant mice.** (**A**, **B**) Representative images show *Gcg* mRNA expression in the PH, detected by FISH, in *ad libitum* fed (**A**) and 48h fasted (**B**) male FVB/Ant mice. (**C**) The graph shows the cumulative FISH signal in PH neurons (integrated density in arbitrary units), measured by image analysis (n=4-5 per group, 9-10 wks). Tissue collection (every 10th 16 µm thick section was collected and analyzed) and imaging parameters were slightly different from the 30h fasting experiment. Mean values are shown above columns. Statistical significance: *** p=0.0003, t-test. *Gcg* expression was as robust in 48h fasted as in 30h fasted FVB/Ant mice. The FISH signal increase was lower (3.5-fold), however, due to the fact that three of the fed mice had noticeably higher *Gcg* expression levels than what we generally observed in other fed FVB/Ant mice. While this could be due to natural variation, in this experiment, mice were housed individually when the experiment began, and this stress and/or its anorexic effect might have contributed to this alteration. Abbreviations: PH, posterior hypothalamic nucleus; pm, principal mammillary tract.

**Figure S3**

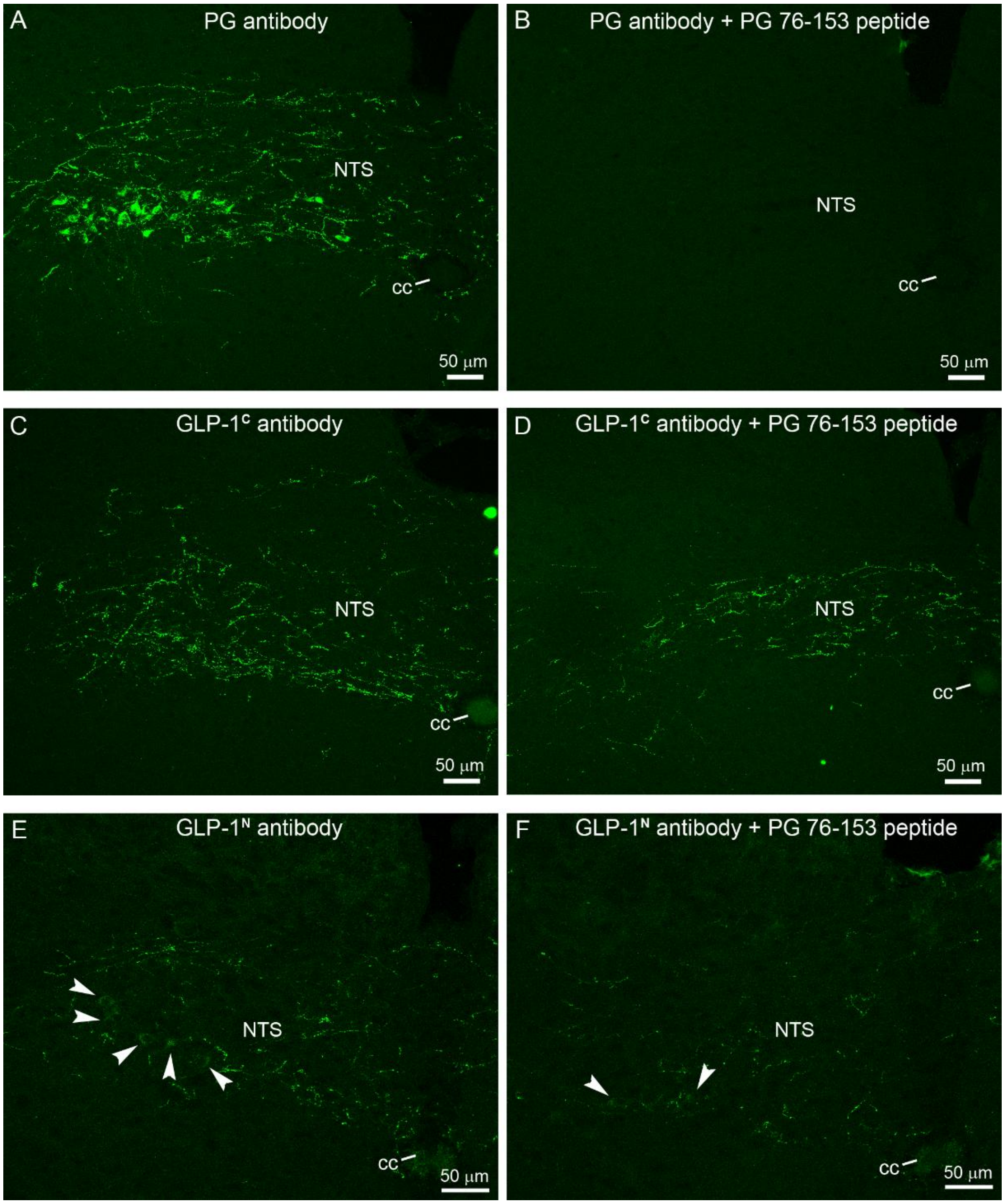


**Figure S3. Antibody preadsorption tests for the PG, GLP-1$^C$ and GLP-1$^N$ antibodies.** The working dilutions of the antibodies were pre-incubated for 3h with a recombinant peptide corresponding to PG 76-153 (human preproglucagon 96-173, ThermoFisher, Cat# RP-102746; 20 µg/ml of the His-ABP-tagged peptide). This peptide contains the entire GLP-1 (7-37) sequence, which corresponds to PG 78-108. Confocal immunofluorescent images show the nucleus of the solitary tract in sections from male FVB/Ant mice. Control sections from the same brains, incubated in the working dilutions of the primary antibodies, were parallel processed. (**A**, **B**) The PG 76-153 peptide completely prevented immunostaining with the PG antibody, which binds to the GLP-1 (12-22) epitope. (**C**, **D**) PG 76-153 did not affect the immunostaining of the GLP-1$^C$ antibody that binds to the amidated C-terminus of GLP-1. (**E**, **F**) In two separate preadsorption tests conducted with the GLP-$1^N$ antibody, which binds the free N-terminus of GLP-1(7-37) or GLP-1(7-36) amide, PG 76-153 either did not affect or slightly reduced the intensity of GLP-$1^N$ immunolabeling. This suggests low-level cross-reactivity with the PG 76-153 peptide, which is N-terminally extended with 2 amino acids compared to GLP-1 (7-37). Arrowheads on **E** and **F** indicate GLP-$1^N$-labeled cell bodies in the NTS. Abbreviations: cc, central canal; NTS, nucleus of the solitary tract.

**Figure S4**

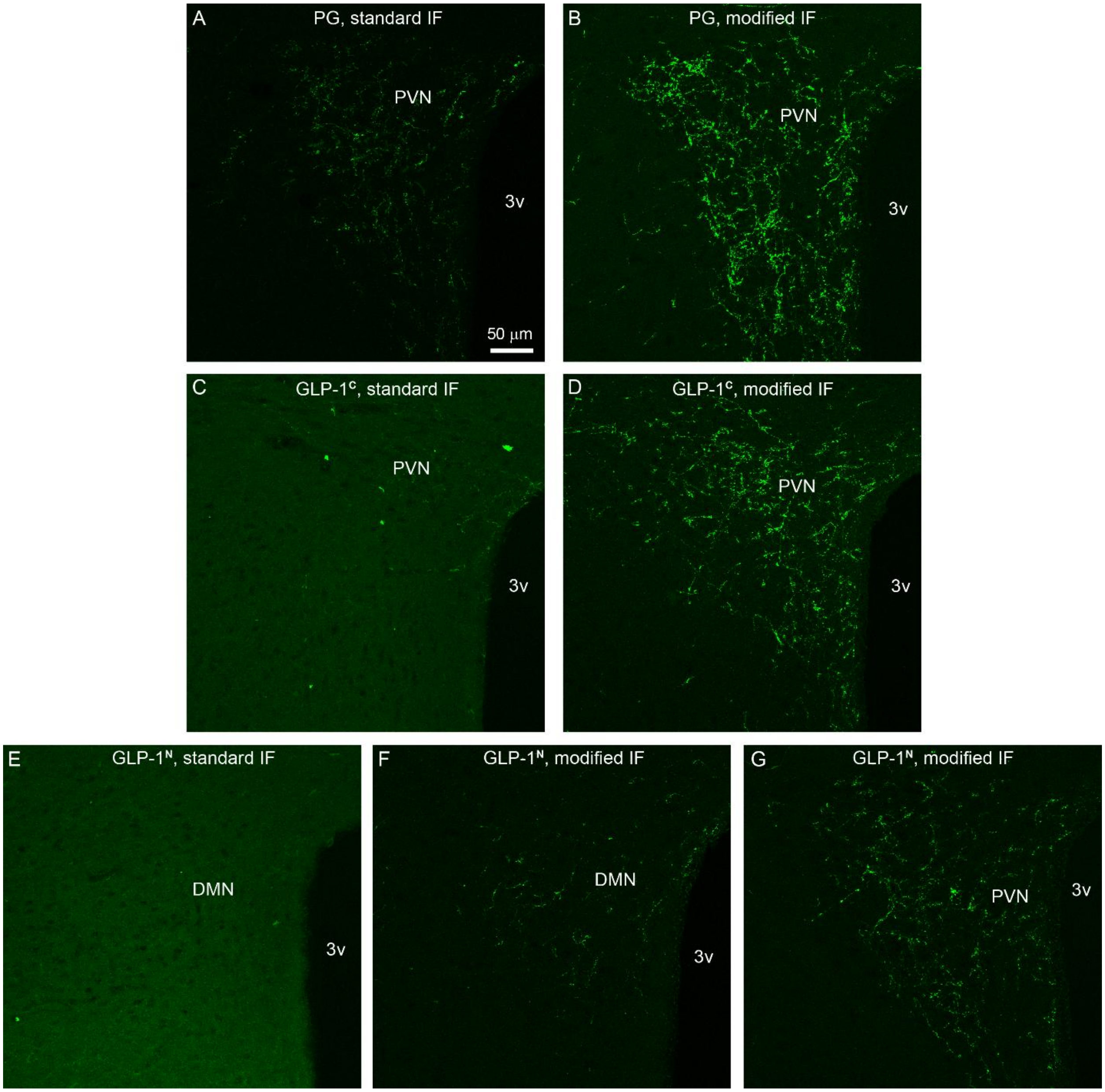


**Figure S4. PG, GLP-1$^{C}$ and GLP-1$^{N}$ IF using the standard and modified IF protocols.** Hypothalamic sections from the same brain (a female C57BL/6J mouse) were parallel processed either through the standard or the modified IF protocol. Primary antibody concentrations were 0.2 µg/ml (PG and GLP-1$^{N}$) or 1 µg/ml (GLP-1$^{C}$). Secondary antibodies were Alexa Fluor 555-conjugated anti-rabbit (for PG) or anti-mouse (for GLP-1$^{C}$ and GLP-1$^{N}$) IgGs. Sections were imaged with a confocal microscope, using the exact same settings, and Z-projections were made from the same number of optical slices. Images show GLP-1-containing axons in the hypothalamic paraventricular and dorsomedial nuclei. (**A**, **B**) PG-labeling is substantially brighter with the modified (**B**) than with the standard protocol (**A**). (**C**, **D**) With the standard protocol, a significant background hindered the detection of specific GLP-1$^{C}$ signal (**C**). The modified protocol both eliminated this background and greatly enhanced GLP-1$^{C}$ labeling (**D**). (**E-G**) GLP-1$^{N}$ labeling was barely discernible using the standard protocol (**E**). The modified protocol eliminated the very high background and allowed detection of specific GLP-$1^N$ labeling (**F**, **G**). Abbreviations: 3v, third ventricle; DMN, dorsomedial nucleus; PVN, paraventricular nucleus.

### Figure S5

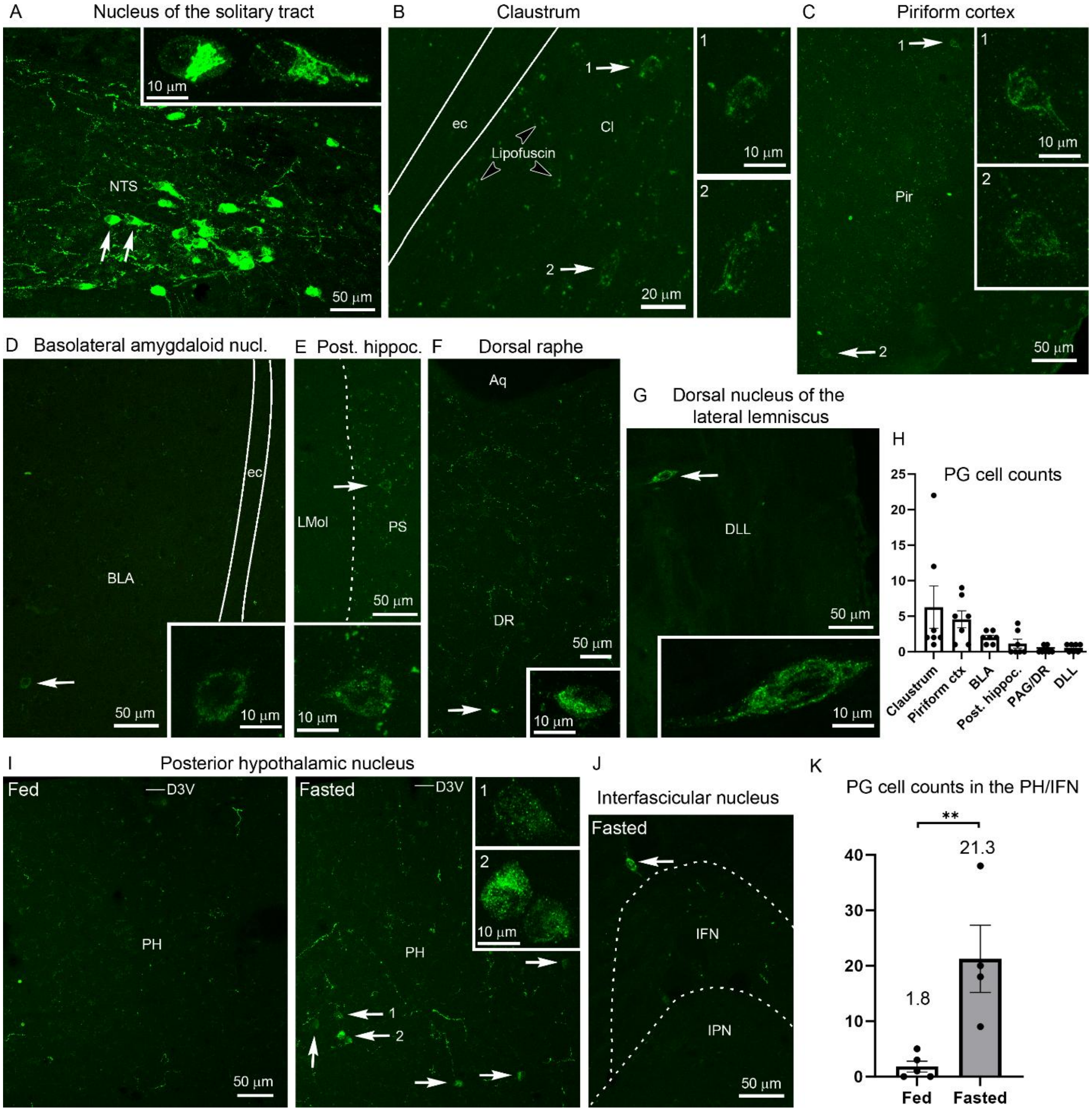


**Figure S5. PG-containing neurons in the brain, detected with the standard IF protocol.** Confocal images of sections from male FVB/Ant mice (11-15 wks). (**A**) PG neurons in the nucleus of the solitary tract were very intensely labeled. The cells indicated by arrows are shown in higher magnification in the inset. (**B-G**) PG neurons (arrows) were less intensely labeled in the claustrum (**B**), piriform cortex (**C**), basolateral amygdaloid nucleus (**D**), posterior hippocampus (**E**), dorsal raphe (**F**), and dorsal nucleus of the lateral lemniscus (**G**). The cells are shown in higher magnification in the insets. The claustrum image (**B**) demonstrates that the intensity of PG-labeling is comparable to the autofluorescence of lipofuscin granules (open arrowheads). Lipofuscin autofluorescence is well-visible in **B**-**F** and **I.** (**H**) Graph showing the number of detected PG neurons through a section series (counted in every 4th 25 µm thick section; n=7). These numbers are a fraction of those detected with the modified IF protocol, shown in Fig S6E. (**I**) Images of the posterior hypothalamic nucleus show no PG neurons in the fed, but several PG neurons in the fasted state (arrows; examples shown in the higher magnification insets). (**J**) A PG neuron (arrow) dorsal to the interfascicular nucleus in a fasted mouse. (**K**) Graph showing the number of PG neurons in the PH/IFN (counted in every 4th 25 μm thick section) in fed and 30h fasted mice (n=4-5 per group; 11-15 wks). Mean values are shown above columns. Statistical significance: ** p=0.0091, t-test. Abbreviations: Aq, aqueduct; BLA, basolateral amygdaloid nucleus; Cl, claustrum; D3V, dorsal third ventricle; DLL, dorsal nucleus of the lateral lemniscus; DR, dorsal raphe; ec, external capsule; IFN, interfascicular nucleus; IPN, interpeduncular nucleus; LMol, lacunosum moleculare layer of the hippocampus; NTS, nucleus of the solitary tract; PH, posterior hypothalamic nucleus; Pir, piriform cortex; PS, prosubiculum.

### Figure S6

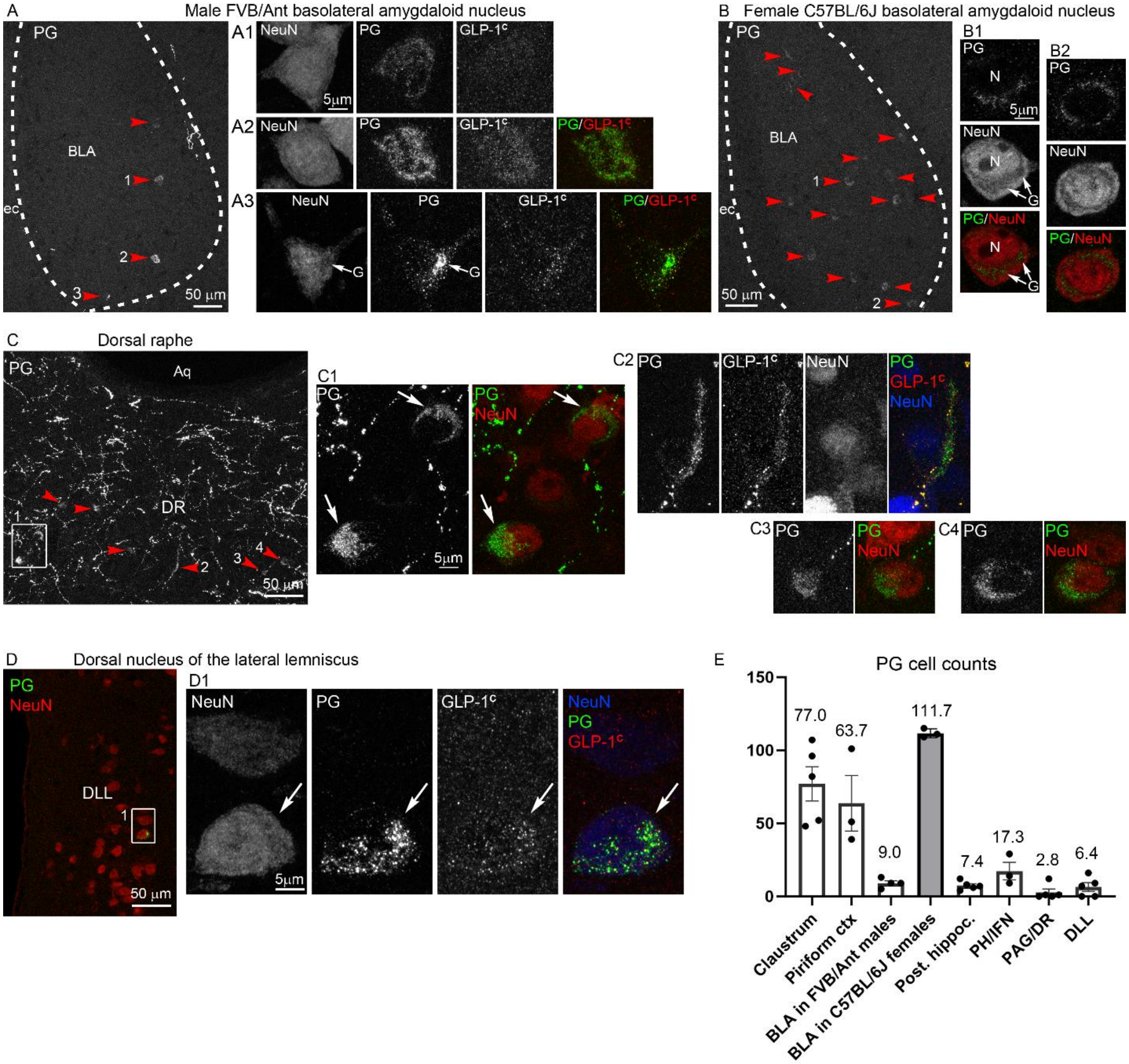


**Figure S6. PG and GLP-1 synthesis in BLA, DR and DLL neurons, and PG cell counts.** (**A**) Triple-label IF for PG/GLP-1$^{C}$ /NeuN in a male FVB/Ant mouse (12 wks). Low magnification image of the PG signal shows four PG neurons (red arrowheads) in the rostral BLA. High magnification confocal images (**A1-A3**) show the NeuN, PG and GLP-1$^{C}$ signals of the cells indicated with numbered arrowheads. **A1** shows a GLP-1$^{C}$-negative, **A2** and **A3** show GLP-1$^{C}$-positive cells (merged PG/GLP-1$^{C}$ panels included). In **A3**, the PG signal labels not only the Golgi apparatus (arrow), but also apparent secretory vesicles in the cell body and dendrites. Many of these vesicles are co-labeled with the GLP-1$^{C}$ antibody. (**B**) The low magnification image shows several PG neurons (red arrowheads) in the BLA of a female C57BL/6J mouse (12 wks). High magnification confocal images (single optical sections) of two cells (**B1, B2**) show that PG is concentrated in the Golgi apparatus (arrows), which appears hollow in the NeuN images. (**C**) PG neurons (red arrowheads, and in the boxed area) in the DR of a male FVB/Ant mouse (15 wks).

High magnification images of the boxed area and numbered cells show the PG and NeuN signals (**C1**-**C4**), as well as the GLP-1$^C$ signal for a GLP-1$^C$ positive cell in **C2**. (**D**) The low magnification image (PG - green/NeuN - red) shows an intensely labeled PG neuron (inside the box) in the DLL of a male FVB/Ant mouse (15 wks). High magnification images of the boxed area show that the PG neuron is positive for GLP-1$^C$ (arrow). (**E**) Graph showing the number of detected PG neurons in 7 PG-expressing neuron population (counted in every 4th 25 μm thick section) in *ad libitum* fed male FVB/Ant mice (n=3-5; 11-15 wks). PG neurons in the BLA were also counted in female C57BL/6J mice (gray column; n=3, 12 wks). Mean values are shown above columns. Abbreviations: Aq, aqueduct; BLA, basolateral amygdaloid nucleus; DLL, dorsal nucleus of the lateral lemniscus; DR, dorsal raphe; ec, external capsule; G, Golgi apparatus.

**Figure S7**

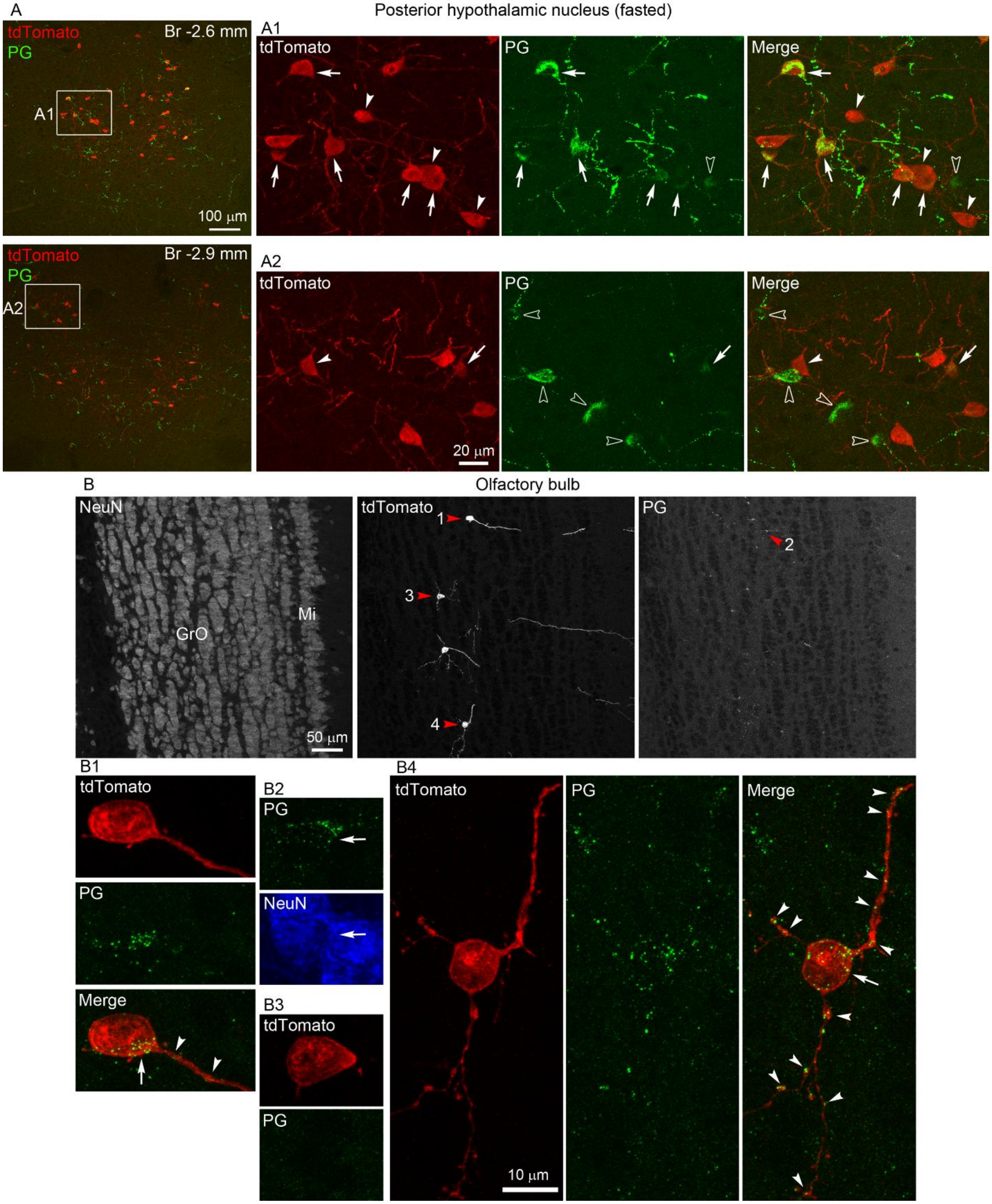


**Figure S7. PG expression in tdTomato neurons of *Gcg*-Cre;tdTomato mice.** (**A**) Dual IF for tdTomato and PG in the posterior hypothalamic nucleus of a 30h fasted female *Gcg-Cre*$^{/+}$*;tdTom*$^{/+}$ mouse. Low magnification merge images (tdTomato - red/PG -green) show sections from two

Bregma levels. Boxed areas are shown in higher magnification confocal images in separate and merge channels (Z-projections). Arrows indicate tdTomato/PG neurons, arrowheads indicate single-labeled tdTomato neurons, open arrowheads indicate single-labeled PG neurons. TdTomato neurons not indicated by an arrow or arrowhead have minimal PG signal. (**B**) Triple IF for NeuN, tdTomato and PG in the olfactory bulb of a *Gcg-Cre*$^{/Cre}$*;tdTom*$^{/tdTom}$ mouse. Cells indicated by red arrowheads are shown in higher magnification. **B1** and **B4** show PG-positive tdTomato cells; PG-containing granules/vesicles are present in both the cell bodies (arrows) and cell processes (arrowheads). **B2** shows a PG cell not expresing tdTomato; the NeuN channel (blue) shows that PG is accumulated in a neuronal cell body (arrow). **B3** shows a PG-negative tdTomato cell. Abbreviations: GrO, granule cell layer of the olfactory bulb; Mi, mitral cell layer of the olfactory bulb.

**Figure S8**

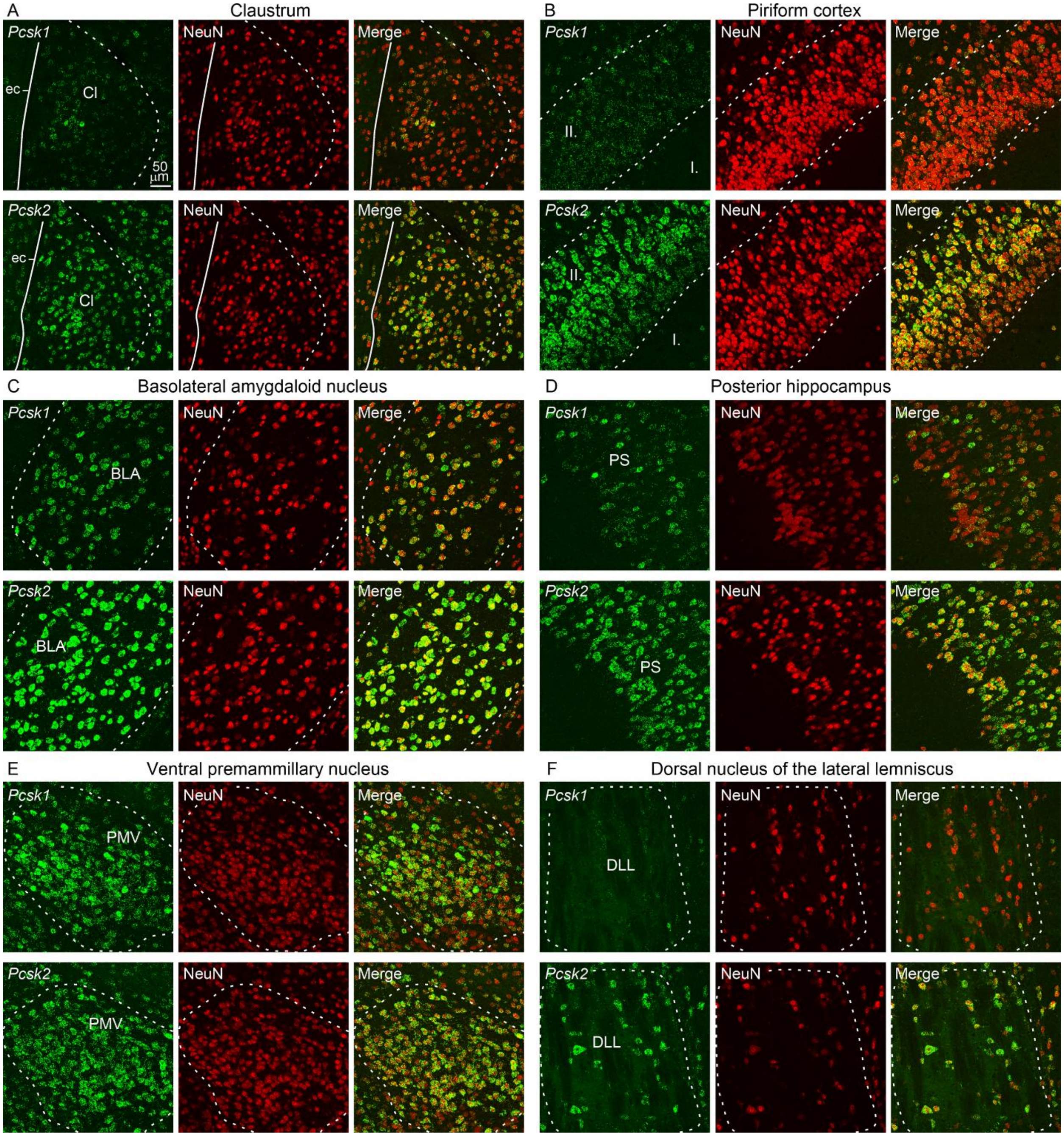


**Figure S8. *Pcsk1* and *Pcsk2* mRNA expression in select *Gcg*-expressing brain regions of C57BL/6J mice.** *Pcsk1* or *Pcsk2* FISH (green) combined with NeuN IF (red) are shown in the (**A**) claustrum, (**B**) piriform cortex, (**C**) rostral part of the basolateral amygdaloid nucleus, (**D**) posterior hippocampus, (**E**) ventral premammillary nucleus, and (**F**) dorsal nucleus of the lateral lemniscus. Adjacent sections were hybridized for *Pcsk1* or *Pcsk2*. In each area either all or the vast majority of neurons express both *Pcsk1* and *Pcsk2* mRNAs. Abbreviations: I.-II., layers I-II. of the piriform cortex; BLA, basolateral amygdaloid nucleus; Cl, claustrum; DLL, dorsal nucleus of the lateral lemniscus; ec, external capsule; PMV, ventral premammillary nucleus; PS, prosubiculum.

**Figure S9**

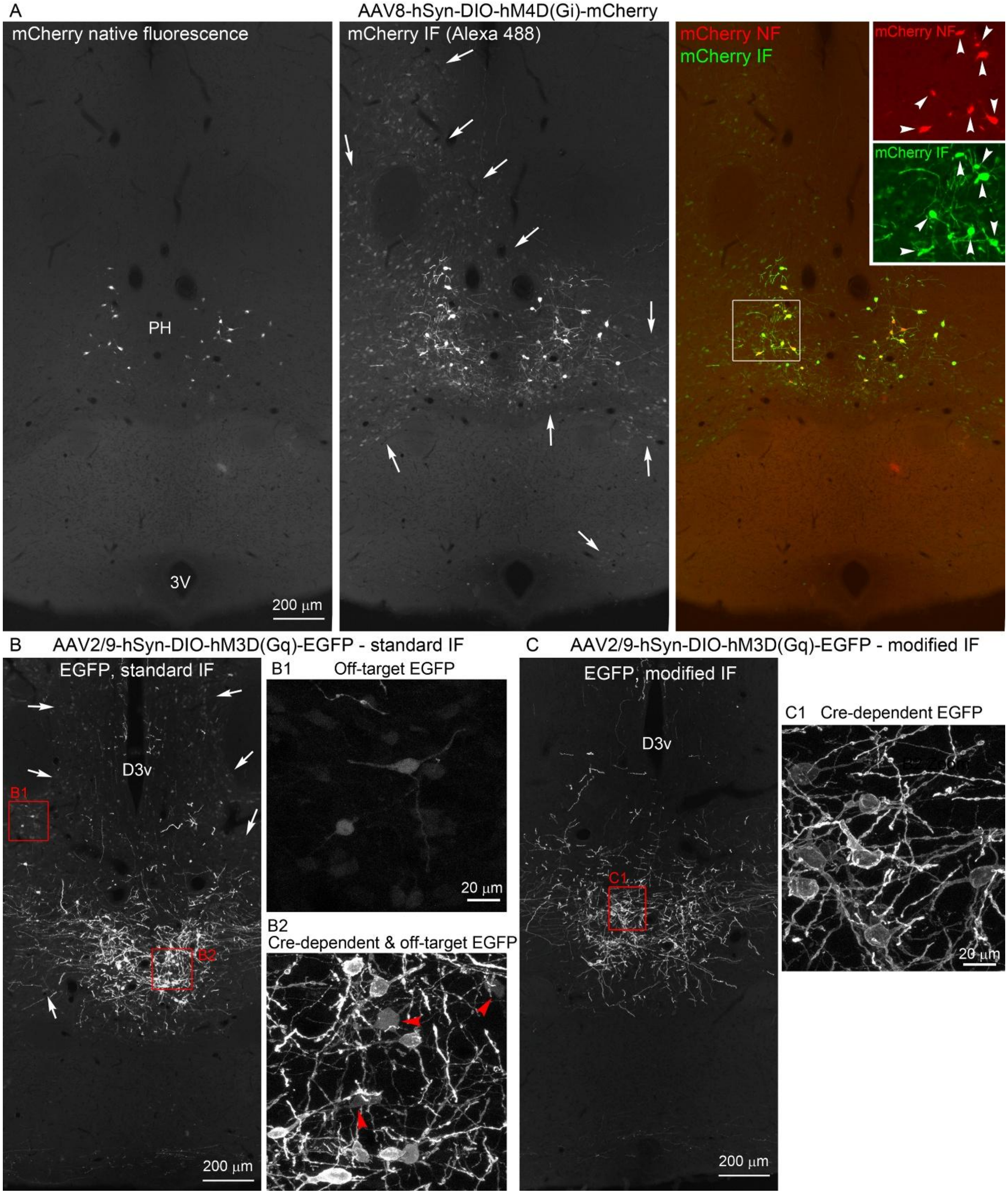


**Figure S9. Off-target expression of Cre-dependent AAVs.** (**A**) mCherry native fluorescence and mCherry IF in a *Gcg*-Cre mouse injected with AAV8-hSyn-DIO-hM4D(Gi)-mCherry in the PH. Left panel: mCherry native fluorescence labels PH neurons in the expected pattern of *Gcg* neurons. Middle panel: mCherry IF (detected with a tdTomato antiserum and Alexa Fluor 488-

conjugated secondary antibody) labels the same cells intensely, and a large number of other neurons near the injection site, less intensely (arrows). Right panel: merged image, the boxed area is magnified to double-size in the insets, showing the separate fluorescent channels. All cells with mCherry native fluorescence are intensely labeled with mCherry IF (arrowheads). Lighter mCherry IF signal is visible in other neurons nearby. (**B-C**) EGFP IF with the standard (**B**) and modifed (**C**) IF protocol in a *Gcg*-Cre mouse injected with AAV2/9-hSyn-DIO-hM3D(Gq)-EGFP into the PH. Sections from the same brain were processed through the respective protocols and labeled with a sheep YPet antiserum and Alexa Fluor 488-conjugated secondary antibody. Boxed areas are shown in higher magnification confocal Z-projection images. (**B**) With standard IF, off-target EGFP expression is detected in a large number of neurons (arrows). **B1**: off-target EGFP expression in neurons, with varying IF signals. **B2**: both Cre-dependent and off-target EGFP expression in neurons. Red arrowheads indicate neurons with off-target EGFP expression, which display less intense and homogenous IF signal. (**C**) The modified IF protocol results in the near-complete elimination of off-target EGFP IF signal. **C1** shows Cre-dependent EGFP expression. The IF signal is most intense along the cell membrane, as EGFP is fused to the transmembrane hM3D(Gq) receptor. Abbreviations: 3V, third ventricle; D3V, dorsal third ventricle; NF, native fluorescence; PH, posterior hypothalamic nucleus.

**Figure S10**

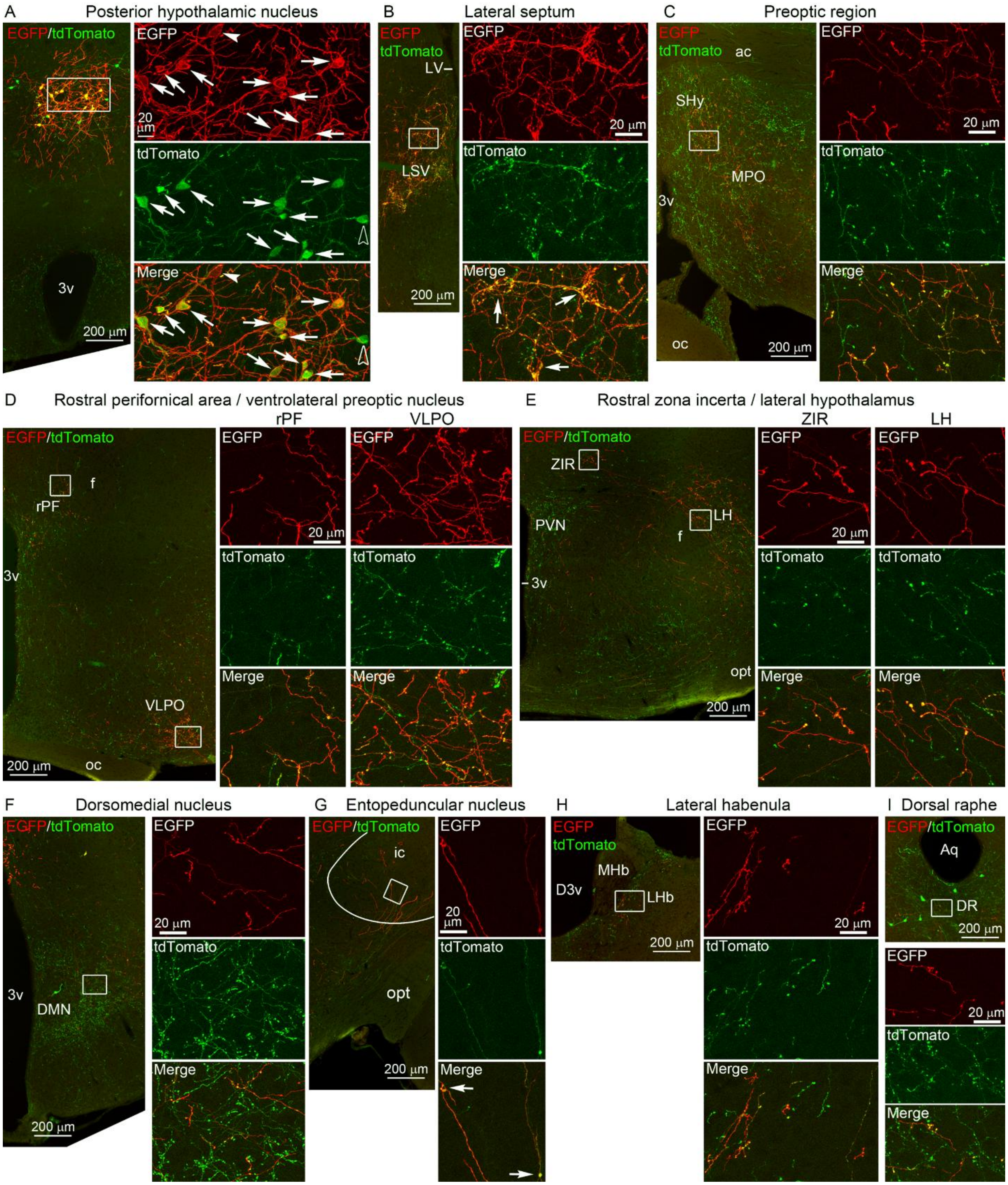


**Figure S10. AAV injections in *Gcg*-Cre;tdTomato mice label tdTomato neurons in the PH/IFN.** Dual IF for EGFP and tdTomato in a male *Gcg-Cre*$^{/+}$;*tdTom*$^{/+}$ mouse injected with AAV2/9-hSyn-DIO-hM3D(Gq)-EGFP in the PH/IFN. To enhance visibility, the more intense EGFP signal is shown in red, and tdTomato in green. The boxed areas are shown in high magnification confocal images (Z-projections), beside the low magnification images. (**A**) In the

PH, most EGFP-labeled neurons express tdTomato and vice versa (arrows), but neurons labeled for only EGFP (arrowhead), or only tdTomato (open arrowhead) are also present. The EGFP signal is associated with the cell membrane since EGFP is fused to the transmembrane receptor hM3d(Gq); tdTomato is expressed in the cytoplasm and nucleus. (**B-I**) In each projection area of PH/IFN *Gcg* neurons, most EGFP-labeled fibers contain tdTomato. EGFP/tdTomato fibers are shown in (**B**) the lateral septum where they form pericellular baskets (arrows), (**C**) preoptic region, (**D**) rostral perifornical area and ventrolateral preoptic nucleus, (**E**) rostral zona incerta and lateral hypothalamus, (**F**) hypothalamic dorsomedial nucleus, (**G**) entopeduncular nucleus inside the internal capsule (arrows point to boutons), (**H**) lateral habenula and (**I**) dorsal raphe. EGFP/tdTomato axons are highly varicose in most areas. Abbreviations: 3v, third ventricle; ac, anterior commissure; Aq, aqueduct; D3v, dorsal third ventricle; DR, dorsal raphe; f, fornix; ic, internal capsule; LH, lateral hypothalamus; LHb, lateral habenula; DMN, hypothalamic dorsomedial nucleus; LSV, ventral lateral septum; LV, lateral ventricle; MHb, medial habenula; MPO, medial preoptic area; oc, optic chiasm; opt, optic tract; rPF, rostral perifornical area; PVN, hypothalamic paraventricular nucleus; SHy, septohypothalamic nucleus; VLPO, ventrolateral preoptic nucleus; ZIR, rostral zona incerta.

**Figure S11**

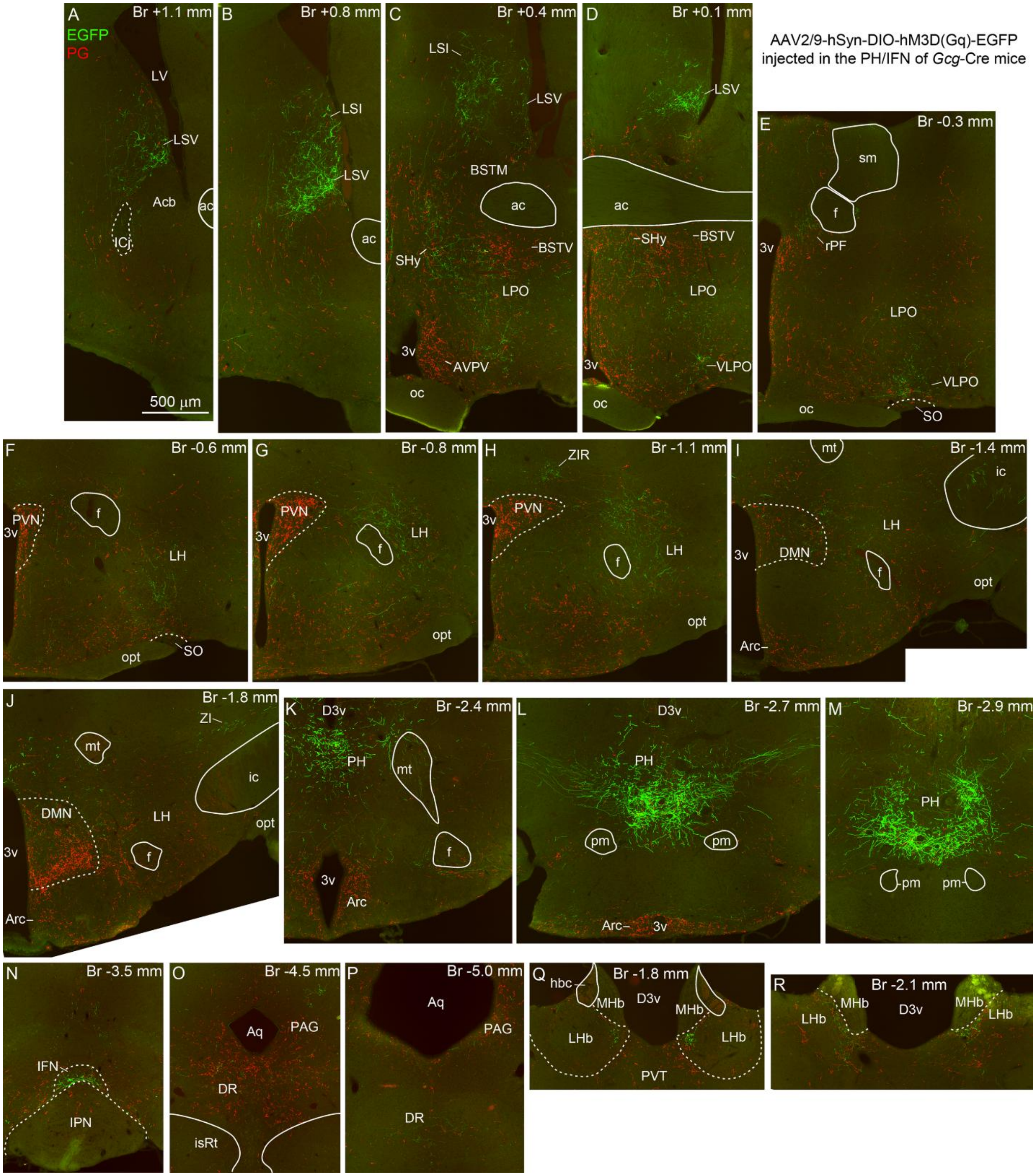


**Figure S11. Projections of PH/IFN and medullary NTS/IRN *Gcg* neurons.** Dual IF for EGFP (green) and PG (red) in a female *Gcg*-Cre[/Cre] mouse injected with AAV2/9-hSyn-DIO-hM3D(Gq)-EGFP in the PH/IFN. This figure shows the same images as Figure 7, with the PG IF signal included. Intensely labeled PG fibers originate from NTS/IRN *Gcg* neurons, EGFP-labeled fibers originate from PH/IFN *Gcg* neurons. Areas receiving dense projections from NTS/IRN, but sparse to none from PH/IFN *Gcg* neurons, include the anteroventral periventricular nucleus (**C**), ventral bed nucleus of the stria terminalis (**C**, **D**), hypothalamic paraventricular nucleus (**F-H**), arcuate nucleus (**I-L**) and paraventricular thalamic nucleus (**Q**). Fibers of NTS/IRN *Gcg* neurons also dominate in the ventral part of the dorsomedial nucleus (**J**) and the PAG/DR (**O**). Abbreviations: 3v, third ventricle; ac, anterior commissure; Acb, nucleus accumbens; Aq, aqueduct; Arc, arcuate nucleus; AVPV, anteroventral periventricular nucleus; BSTM, medial part of the bed nucleus of the stria terminalis; BSTV, ventral part of the bed nucleus of the stria terminalis; D3v, dorsal third ventricle; DR, dorsal raphe; f, fornix; hbc, habenular commissure; ic, internal capsule; ICj, island of Calleja; IFN, interfascicular nucleus; IPN, interpeduncular nucleus; isRt, isthmic reticular formation; LH, lateral hypothalamus; LHb, lateral habenula; DMN, hypothalamic dorsomedial nucleus; LPO, lateral preoptic area; LSI, intermediate lateral septum; LSV, ventral lateral septum; LV, lateral ventricle; MHb, medial habenula; mt, mammillothalamic tract; oc, optic chiasm; opt, optic tract; PAG, periaqueductal gray; PH, posterior hypothalamic nucleus; pm, principal mammillary tract; rPF, rostral perifornical area; PVN, hypothalamic paraventricular nucleus; PVT, thalamic paraventricular nucleus; SHy, septohypothalamic nucleus; sm, stria medullaris; SO, supraoptic nucleus; VLPO, ventrolateral preoptic nucleus; ZI, zona incerta; ZIR, rostral zona incerta.

**Figure S12**

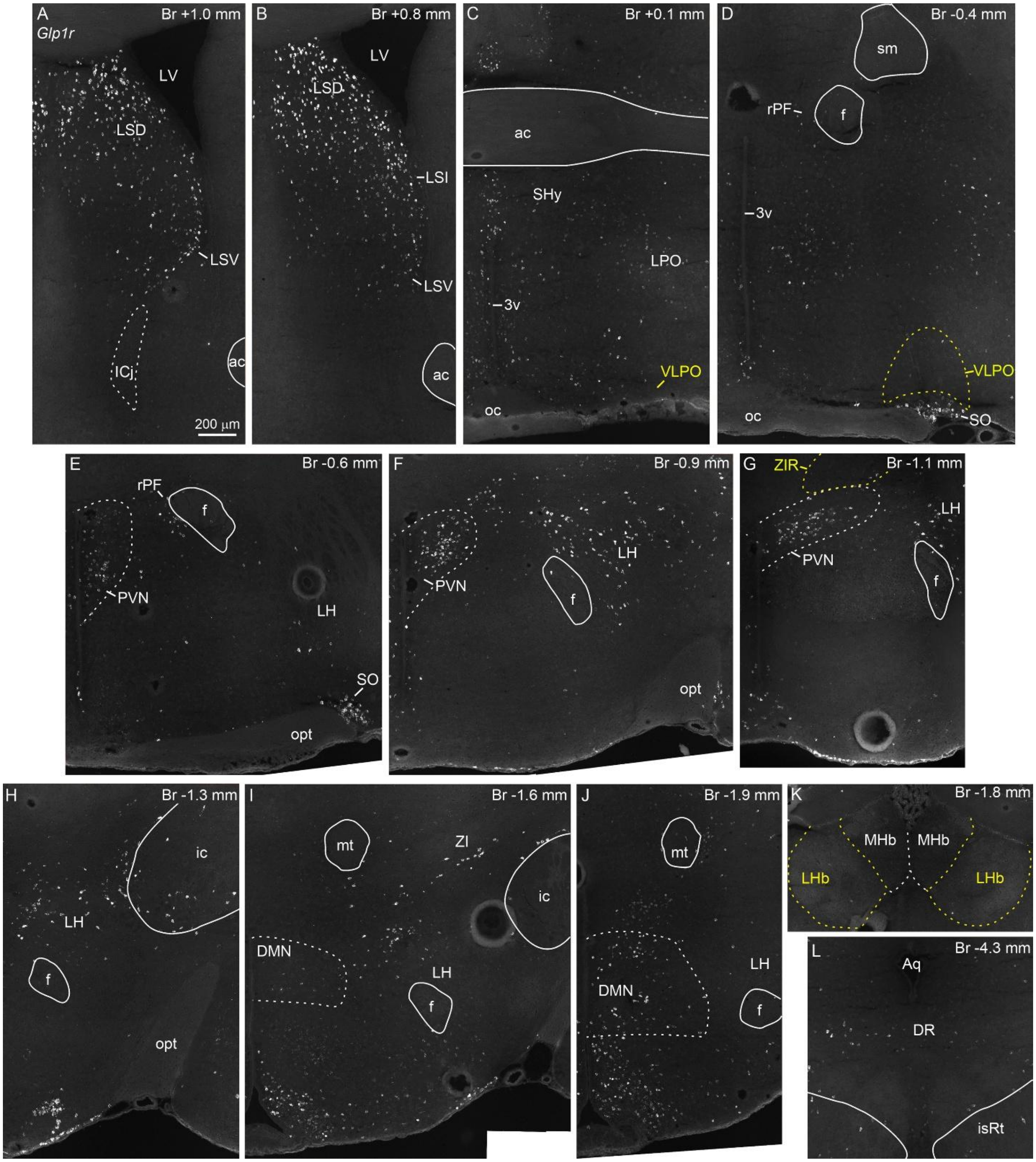


**Figure S12.** ***Glp1r* mRNA expression in the projection areas of PH/IFN *Gcg* neurons.** FISH for *Glp1r* mRNA in a male C57BL/6J mouse. *Glp1r* is abundantly expressed in most areas where PH/IFN Gcg neurons project, including the ventral and intermediate lateral septum (**A**, **B**), preoptic region (**C**), rostral perifornical area (**D**, **E**), lateral hypothalamus (**E-I**), entopeduncular nucleus inside the internal capsule (**H**), dorsomedial nucleus (**I**, **J**), lateral zona incerta (**I**) and dorsal raphe (**L**). *Glp1r*-expressing neurons are very rare in the ventrolateral preoptic nucleus (**C**, **D**), rostral zona incerta (**G**) and lateral habenula (**K**). Abbreviations: 3v, third ventricle; ac, anterior commissure; Aq, aqueduct; DR, dorsal raphe; f, fornix; ic, internal capsule; ICj, island of Calleja; isRt, isthmic reticular formation; LH, lateral hypothalamus; LHb, lateral habenula; DMN, hypothalamic dorsomedial nucleus; LPO, lateral preoptic area; LSD, dorsal lateral septum; LSI, intermediate lateral septum; LSV, ventral lateral septum; LV, lateral ventricle; MHb, medial habenula; mt, mammillothalamic tract; oc, optic chiasm; opt, optic tract; rPF, rostral perifornical area; PVN, hypothalamic paraventricular nucleus; sm, stria medullaris; SHy, septohypothalamic nucleus; SO, supraoptic nucleus; VLPO, ventrolateral preoptic nucleus; ZI, zona incerta; ZIR, rostral zona incerta.

**Supplementary Table 1. TdTomato cell counts in *Gcg*-Cre;tdTomato mice.** Neurons with native tdTomato fluorescence were counted bilaterally on every 4th 25 µm thick section through a neuron population, or the entire brain. Amy, Amygdala region; C, cerebrum; Cb, Cerebellum; Hy, Hypothalamus; Mi, Midbrain; P-M, Pons-Medulla; Th, Thalamus; ZI, Zona incerta.
Tdtomato neurons were not counted through the OB, but *Gcg-Cre*$^{/\mathrm{Cre}}$;*tdTom*$^{/tdTom}$ brains had approximately twice as many tdTomato neurons in the OB than *Gcg-Cre*$^{/+}$;*tdTom*$^{/+}$ brains (up to 40 *vs* up to 25 per section). * Pre-fasted: mice were fasted for 24h, twelve days before euthanasia.

| | Age | Feeding | PH/IFN | PAG/ | | NTS/ | | | Other areas in the | | | | | |
|---|---|---|---|---|---|---|---|---|---|---|---|---|---|---|
| Sex | (days) | status | (PH+IFN) | DR | DLL | IRN | Amy | ZI | C | Th | Hy | Mi | P-M | Cb |
| *Gcg-Cre*$^{/+}$;*tdTom*$^{/+}$ | | | | | | | | | | | | | | |
| M | 124 | *Ad Lib* | 80 (73+7) | 59 | 15 | 328 | 66 | 30 | 32 | 3 | 8 | 5 | 8 | 3 |
| M | 124 | *Ad Lib* | 55 (53+2) | 44 | 7 | 290 | n/a | 27 | n/a | n/a | 17 | 4 | n/a | n/a |
| M | 66 | *Ad Lib* | 67 (65+2) | 23 | 3 | 319 | 40 | 15 | 25 | 0 | 1 | 6 | 3 | 4 |
| M | 66 | *Ad Lib* | 56 (53+3) | 17 | 7 | n/a | | | | | | | | |
| M | 66 | Pre-fasted* | 124 (110+14) | n/a | n/a | n/a | | | | | | | | |
| M | 66 | Pre-fasted* | 83 (83+0) | 18 | 7 | 342 | | | | | | | | |
| F | 88 | *Ad Lib* | 111 (101+10) | 30 | 7 | 373 | | | | | | | | |
| F | 77 | *Ad Lib* | 119 (111+8) | 20 | 15 | 298 | | | | | | | | |
| F | 76 | *Ad Lib* | 83 (78+5) | 19 | 7 | 326 | | | | | | | | |
| F | 66 | *Ad Lib* | 122 (114+8) | 21 | 9 | n/a | | | | | | | | |
| *Gcg-Cre*$^{/\mathrm{Cre}}$;*tdTom*$^{/tdTom}$ | | | | | | | | | | | | | | |
| F | 61 | *Ad Lib* | 226 (189+37) | 78 | 45 | 391 | 205 | 99 | 108 | 31 | 29 | 37 | 33 | 19 |
| F | 61 | *Ad Lib* | 182 (171+11) | 65 | 46 | n/a | 181 | 93 | 138 | 15 | 30 | 35 | 20 | 16 |
| F | 142 | *Ad Lib* | 294 (263+31) | 117 | 54 | n/a | | | | | | | | |

**Supplementary Table 2. List of AAV-injections into the PH/IFN in *Gcg*-Cre mice.**
AAV-encoded fluorescent reporter (mCherry, EYFP or EGFP)-positive neurons were counted on every 4th 25 µm thick section through the PH and midbrain. Only neurons with intense IF signal were counted. Scattered reporter-positive neurons were also observed dorsal to the PH or IFN. Fasting: mice were fasted ~1-2 weeks after AAV injection for 24h, or before transcardial perfusion for 30h. Antibodies: the tdTomato antibody was used to detect mCherry, the YPet antibody to detect EYFP and EGFP. Abbreviations: std IF, standard immunofluorescence protocol; mod IF, modified immunofluorescence protocol. Cli, caudal linear nucleus of the raphe; RLi, rostral linear nucleus of the raphe.

| Sex | Genotype | AAV | Fasting | Antibodies used for IF | Fluorescent reporter-positive neurons in | | | | | |
|---|---|---|---|---|---|---|---|---|---|---|
| | | | | | PH | IFN | Dorsal to PH+IFN | RLi/ CLi | PAG | DR |
| M | $Cre^{/+}$ | AAV1-EF1a-DIO-hM3D-mCherry | 24h | tdTomato (std IF) | 105 | 8 | 3+0 | 0 | 0 | 0 |
| M | $Cre^{/+}$ | AAV1-EF1a-DIO-hM3D-mCherry | 24h | tdTomato (std IF) | 77 | 3 | 0+1 | 3 | 0 | 0 |
| F | $Cre^{/Cre}$ | AAV8-EF1a-DIO-hM3D-mCherry | - | tdTomato (std IF) | 106 | 14 | 1+4 | 0 | 2 | 0 |
| F | $Cre^{/Cre}$ | AAV8-hSyn-DIO-hM3D-mCherry | - | tdTomato (std IF) | 85 | 28 | 0+1 | 0 | 0 | 0 |
| M | $Cre^{/+}$ | AAV8-hSyn-DIO-hM4D-mCherry | 24h | tdTomato (std IF) | 125 | 8 | 3+0 | 0 | 0 | 0 |
| F | $Cre^{/Cre}$ | AAV2/9-hSyn-DIO-hM4D-EYFP | - | YPet/GLP-$1^C$/PG (mod IF) | 64 | 14 | 7+6 | 5 | 27 | 25 |
| F | $Cre^{/Cre}$ | AAV2/9-hSyn-DIO-hM4D-EYFP | - | YPet (std IF) | 67 | 0 | 1+0 | 0 | 0 | 0 |
| F | $Cre^{/Cre}$ | AAV2/9-hSyn-DIO-hM4D-EYFP | 24h+30h | YPet/PG/GLP-$1^C$ (mod IF) | 61 | 0 | 4+0 | 0 | 5 | 0 |
| F | $Cre^{/Cre}$ | AAV2/9-hSyn-DIO-hM3D-EGFP | 24h+30h | YPet/PG/GLP-$1^C$ (mod IF) | 88 | 15 | 9+2 | 5 | 36 | 58 |
| F | $Cre^{/Cre}$ | AAV2/9-hSyn-DIO-hM3D-EGFP | 24h | YPet/GLP-1R/NeuN (std IF) | 113 | 23 | 22+3 | 1 | 1 | 0 |
| F | $Cre^{/Cre}$ | AAV2/9-hSyn-DIO-hM3D-EGFP | 24h | YPet/PG (mod IF)<br>YPet/GLP-1R/NeuN (std IF) | 93 | 15 | 14+0 | 3 | 8 | 0 |

**Supplementary Table 3.** Antibodies tried for proglucagon detection

| Epitope/immunogen | Source | Antibody Code | Type | Results |
|---|---|---|---|---|
| GLP-1 (12-22) | Novo Nordisk | 1212-0000-0542-1b; (Clone 62-2F6) | Rabbitized monoclonal | Specific, strongest signal |
| Glucagon, midportion | Novo Nordisk | 1212-0000-0554-1b | Rabbitized monoclonal | Specific, good signal with modified IF protocol |
| GLP-1 (7-37) | BMA Biomedicals | T-4363 | Rabbit polyclonal | Specific, good signal |
| Glicentin | ThermoFisher | PA5-89937 | Rabbit polyclonal | Specific, good signal with modified IF protocol |
| Proglucagon (61-110) | Sigma Aldrich | SAB4501137 | Rabbit polyclonal | Neuronal background with both protocols |
